# Hippocampo-neocortical interaction as compressive retrieval-augmented generation

**DOI:** 10.1101/2024.11.04.621950

**Authors:** Eleanor Spens, Neil Burgess

**Affiliations:** Institute of Cognitive Neuroscience, University College London

## Abstract

Many aspects of learning, memory, and problem solving involve interplay between episodic (hippocampal) and semantic (neocortical) systems, but the neural mechanisms supporting this are unclear. We present a computational model in which sequential experiences are encoded in hippocampus in compressed form and replayed to train a neocortical generative network. This network captures the gist of specific episodes and extracts statistical patterns that generalise to new situations, enabling efficient reconstruction of the past and prediction of the future. The two systems interact during encoding, recall and problem solving, with the hippocampus retrieving relevant episodic information into working memory as a basis for generation using the ‘general knowledge’ of the neocortical network. We simulate this interaction as ‘retrieval-augmented generation’, with the addition of mechanisms to *compress* episodic memories into hippocampus and to *consolidate* them into neocortex. The model explains changes to memories over time, including schema-based distortions, and shows how episodic and semantic memory contribute to problem solving.

## 1 Introduction

Despite the success of the theorised distinction between episodic (hippocampal) and semantic (neocortical) systems, and long-standing ideas regarding the consolidation of memories from hippocampus to neocortex (*1–4*), how both systems work together to support a wide variety of cognitive functions is unclear. These include recollection and problem solving, which rely on a variable mixture of specific episodic memories from the hippocampus *and* more general knowledge from the neocortex (*5, 6*). Here we lay out a framework for how this co-operation occurs, and make use of recent advances in machine learning to simulate how the resulting system would perform these functions for comparison with human performance. This builds on many years of theoretical insights into the relationship between schemas and memory (*7–12*), and their potential correspondence to large language models (*13, 14*).

We explore three questions, and implement our hypothesised solutions as a computational model (see Table 1):

1. *How does episodic memory influence semantic memory?* We hypothesise that a deep generative model in neocortex is trained to predict its own inputs, using memories of episodic experiences replayed from hippocampus (and during the experiences themselves). In this way it develops generalisable (semantic) knowledge, enabling it to generate episodes from an initial input. With enough consolidation, neocortex can approximately reconstruct memories without the hippocampus, allowing the redundant hippocampal trace to fade away.
2. *How does semantic memory influence episodic memory?* We hypothesise that the hippocampus, which must use neocortical representations for input and output, stores episodic memories in a highly compressed form, as a minimal subset of information from which the generative model can reconstruct each episode, combining both general schemas and episode-unique features.
3. *How do semantic and episodic memory interact during recall and problem solving?* We hypothesise that hippocampus and neocortex work together to solve problems, with the hippocampus providing specific episodes and the neocortex generating predictions from semantic memory informed by these episodes. In other words, episodic contributions to problem solving correspond to the hippocampus retrieving relevant memories on which to base neocortical generation in working memory. Simply recalling a memory also requires neocortical predictions to reconstruct an event from its compressed hippocampal trace.

**Table 1:**
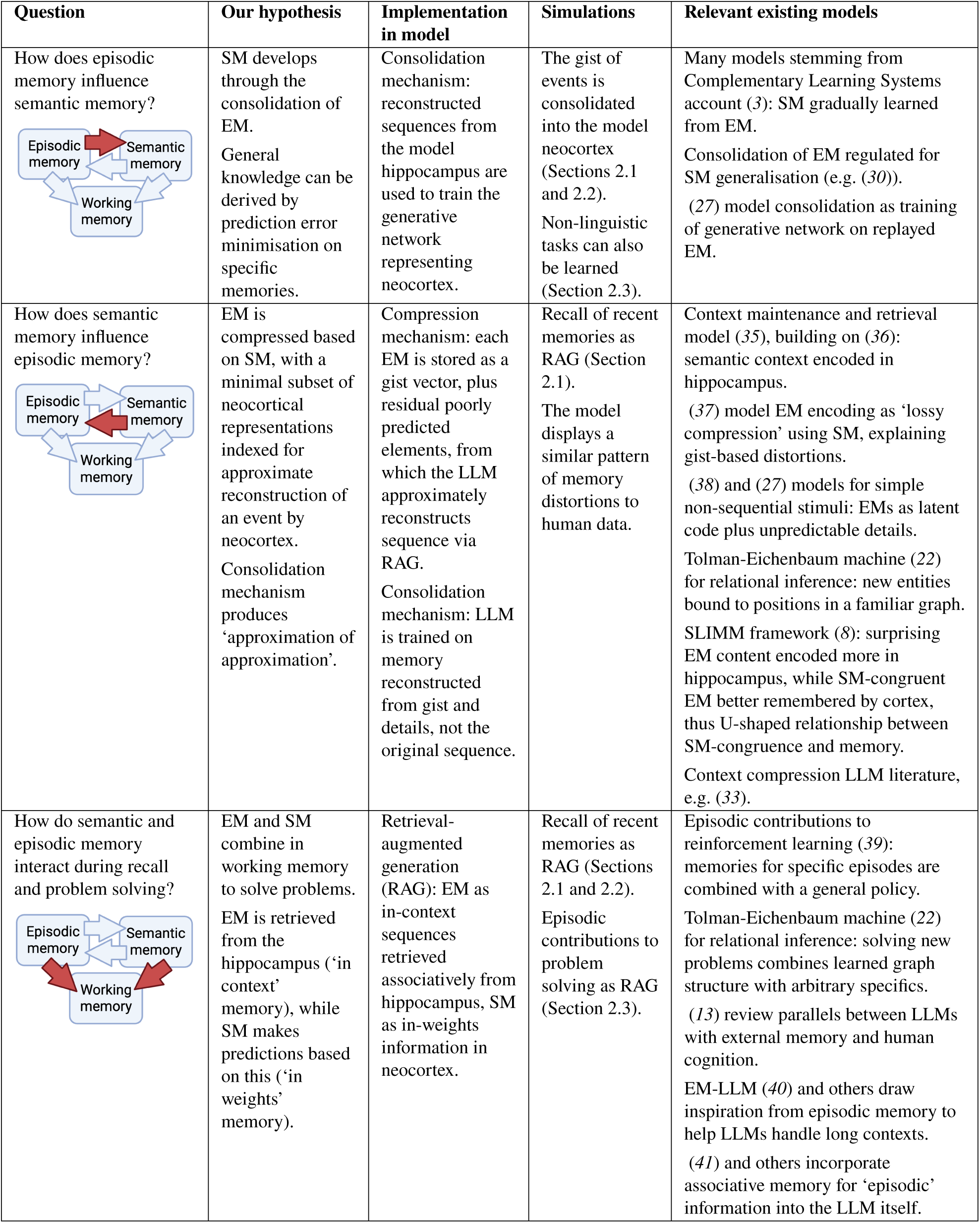
Three questions about the interaction between episodic memory (EM), semantic memory (SM), and working memory (WM), our hypothesised solutions, how we implement our hypotheses computationally using large language models (LLMs) and retrieval-augmented generation (RAG), the relevant results, and relevant previous models.

| Question | Our hypothesis | Implementation in model | Simulations | Relevant existing models |
| --- | --- | --- | --- | --- |
| How does episodic memory influence semantic memory? | SM develops through the consolidation of EM.<br>General knowledge can be derived by prediction error minimisation on specific memories. | Consolidation mechanism: reconstructed sequences from the model hippocampus are used to train the generative network representing neocortex. | The gist of events is consolidated into the model neocortex (Sections 2.1 and 2.2).<br>Non-linguistic tasks can also be learned (Section 2.3). | Many models stemming from Complementary Learning Systems account (3): SM gradually learned from EM.<br>Consolidation of EM regulated for SM generalisation (e.g. (30)).<br>(27) model consolidation as training of generative network on replayed EM. |
| How does semantic memory influence episodic memory? | EM is compressed based on SM, with a minimal subset of neocortical representations indexed for approximate reconstruction of an event by neocortex.<br>Consolidation mechanism produces 'approximation of approximation'. | Compression mechanism: each EM is stored as a gist vector, plus residual poorly predicted elements, from which the LLM approximately reconstructs sequence via RAG.<br>Consolidation mechanism: LLM is trained on memory reconstructed from gist and details, not the original sequence. | Recall of recent memories as RAG (Section 2.1).<br>The model displays a similar pattern of memory distortions to human data. | Context maintenance and retrieval model (35), building on (36): semantic context encoded in hippocampus.<br>(37) model EM encoding as 'lossy compression' using SM, explaining gist-based distortions.<br>(38) and (27) models for simple non-sequential stimuli: EMs as latent code plus unpredictable details.<br>Tolman-Eichenbaum machine (22) for relational inference: new entities bound to positions in a familiar graph.<br>SLIMM framework (8): surprising EM content encoded more in hippocampus, while SM-congruent EM better remembered by cortex, thus U-shaped relationship between SM-congruence and memory.<br>Context compression LLM literature, e.g. (33). |
| How do semantic and episodic memory interact during recall and problem solving? | EM and SM combine in working memory to solve problems.<br>EM is retrieved from the hippocampus ('in context' memory), while SM makes predictions based on this ('in weights' memory). | Retrieval-augmented generation (RAG): EM as in-context sequences retrieved associatively from hippocampus, SM as in-weights information in neocortex. | Recall of recent memories as RAG (Sections 2.1 and 2.2).<br>Episodic contributions to problem solving as RAG (Section 2.3). | Episodic contributions to reinforcement learning (39): memories for specific episodes are combined with a general policy.<br>Tolman-Eichenbaum machine (22) for relational inference: solving new problems combines learned graph structure with arbitrary specifics.<br>(13) review parallels between LLMs with external memory and human cognition.<br>EM-LLM (40) and others draw inspiration from episodic memory to help LLMs handle long contexts.<br>(41) and others incorporate associative memory for 'episodic' information into the LLM itself. |

We implement these ideas in a computational model in which compressed hippocampal memories relevant to a task are retrieved into working memory as context for a neocortical generative network, which is trained on replayed hippocampal memories through consolidation. This enables the generative network to develop generalisable semantic knowledge which supports problem solving, to reconstruct episodes efficiently from minimal hippocampal traces, and eventually to capture the facts of specific events. Our model reflects the complex interplay between episodic and semantic memory; semantic memory is acquired by learning to reconstruct episodic memories, but also shapes the representation of these episodes, both in terms of their initial encoding and their gradual schematisation.

Here working memory refers to the temporary retention of a limited amount of information (*15*), including episodic memories retrieved from long-term memory into the ‘episodic buffer’ (*16*), which can accommodate considerably more than the 4 or so items that can be attended to at a given moment (*16, 17*). This is the ‘workspace’ into which episodic memories relevant to a task are retrieved from the hippocampus, upon which the neocortical network operates.

We first consider simple narratives representing autobiographical episodes, and demonstrate how they are i) encoded in compressed form in hippocampus, ii) retrieved and decompressed by the neocortical generative model, and iii) consolidated into the neocortical generative model. We show that the model predicts changes to the content of memory over time, such as greater abstraction and the loss of specific detail, as seen in human memory (*18, 19*). We also show that the compression aspect of the model enables efficient storage and reconstruction, suggesting how rich and complex events can be stored over a lifetime without exceeding limited hippocampal capacity. Next, we show how encoding and consolidation don’t just result in loss of detail, but systematically distort memories towards expectations in our model, matching experimental data (*20, 21*). Finally, we show how consolidation develops generalisable knowledge in the neocortical network (*22*), allowing problem solving by combining this with relevant specifics, which are loaded into working memory from hippocampus or ongoing experience. Our model works with both linguistic and non-linguistic stimuli, and is thus applicable to sequential experience of many kinds.

### Implementing the model via compressive retrieval-augmented generation

Here we use transformer-based large language models (*23*) to simulate how neocortical networks combine with the hippocampus to extract semantic knowledge from sequential experiences, and to support memory and problem solving.

Machine learning has shown how deep neural networks can act as task-general ‘foundation models’ (*24*). In particular, large language models (LLMs) demonstrate that complex sequential behaviours can develop as a byproduct of a simple ‘next item prediction’ task (*23, 25*), with distributed semantic representations slowly learned over time, in a way analogous to the development of semantic memory in neocortex (*26*). (See Section 4.1 for further details.) Here we suggest that memory consolidation corresponds to the development of neural ‘foundation models’ in a similar way, with neocortex learning to reconstruct replayed memories via schemas (see (*27*)). Crucially, LLMs can memorise specific sequences as well as learning generalities (*28*), meaning that both event-unique information and patterns across events can be captured within the same network. We model consolidation as the training of autoregressive sequence models, simulated using Mistral 7B (*29*) and GPT-2 (*23*), on sequences which represent replayed hippocampal memories. Consolidation can thus be thought of as teacher-student learning (see (*30*)). The objective during training is simply to predict the next item in sequences from the training data. Once trained, the network can continue an input sequence, or generate a new sequence from scratch, by iteratively predicting the next item from the items in the ‘prompt’ or ‘context’ which models the content of working memory.

The hippocampus is modelled simply using modern Hopfield networks (*31*), storing states which can be retrieved autoassociatively as well as the one-step transitions between them. Whilst not the focus of this paper, this illustrates the hypothesis that hippocampus and neocortex use very different mechanisms for sequence memory, though both bind together neocortical representations.

We propose that the hippocampal ‘memory bank’ and neocortical generative network work together to enable problem solving, with the hippocampus retrieving memories relevant to a new task into working memory as context for the generative network. Specifically, we draw inspiration from retrieval-augmented generation (RAG; (*32*)), which involves prompting LLMs with relevant data retrieved from external memory, as a model of drawing on relevant memories to inform predictions in a new situation (see also (*13*)). A typical task for RAG might be question answering based on a set of new facts which do not feature in the training data for the LLM. To ask the system a question, first the most relevant items are found in the external memory (using some similarity metric). A ‘prompt’ is then constructed, in which the retrieved items are given to the LLM together with the question (see Figure 1).

**Figure 1:**
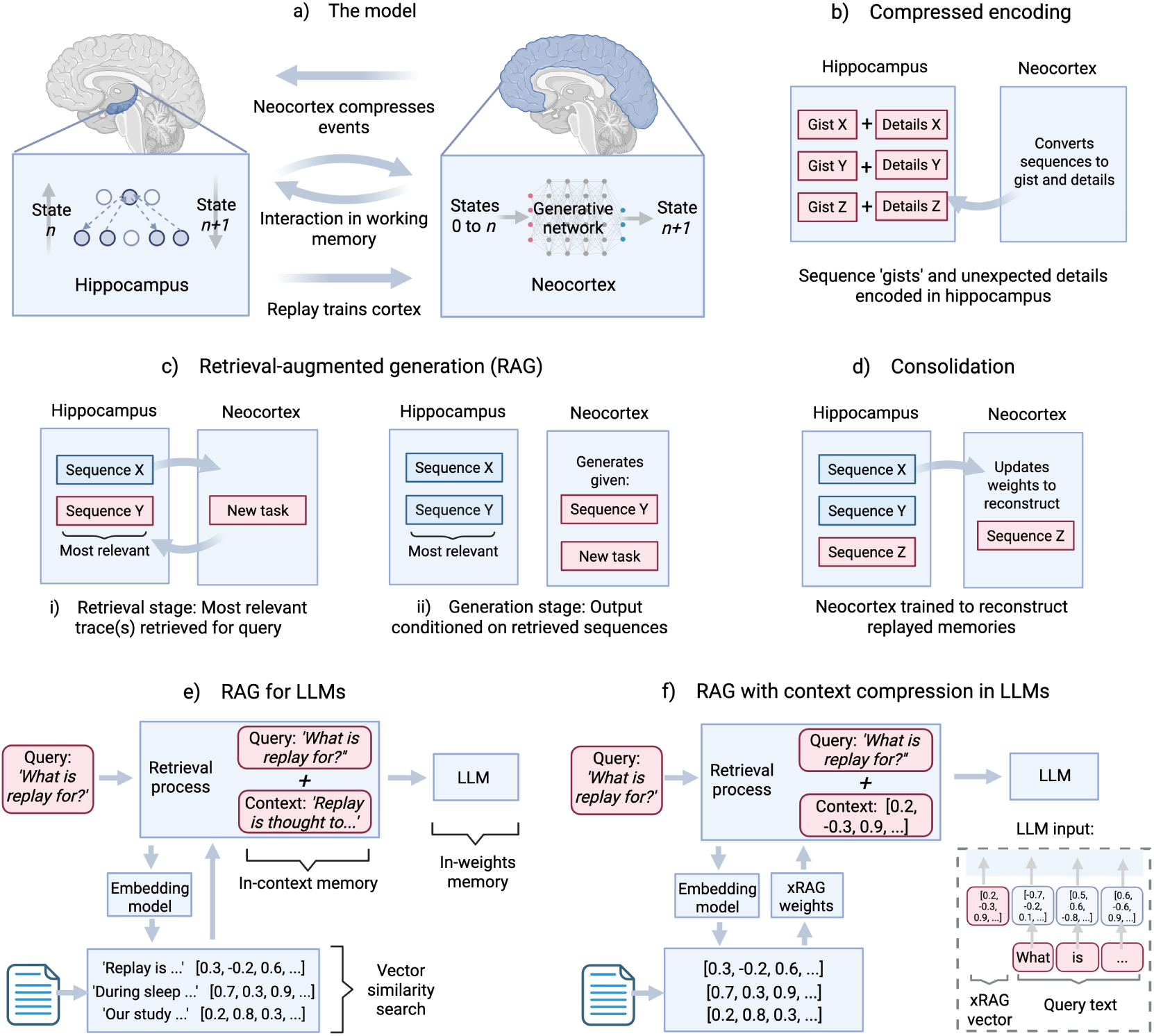
Summary of model. a) Episodic memories are first encoded in the hippocampus, then replayed during rest to train a generative network (simulated by an LLM) which thereby acquires semantic knowledge. The hippocampus and neocortex interact to play complementary roles during problem solving, resembling ‘retrieval-augmented generation’ (RAG; (*32*)). Relevant hippocampal traces are retrieved into working memory and used to ‘prompt’ the neocortical network. The hippocampus is modelled with modern Hopfield networks (*31*) storing states and one-step transitions between them. b) Sequences are encoded in the hippocampus in compressed form. Specifically, a vector from which the sequence can be approximately decoded by neocortex is stored for each event, together with the subsequences that are most surprising given this compressed version of the event. c) The neocortical and hippocampal networks work together to recall memories and solve problems via RAG, e.g. when encountering a new task which requires inferences to be drawn based on hippocampal memories. d) Memories are replayed from the hippocampus during rest, and the generative network captures their statistics through prediction error minimisation. e) RAG for LLMs. When the generative network (neocortex) is prompted with a query, related information is retrieved from external memory (hippocampus). The retrieval process finds stored information with similar vectors (produced by a ‘text embedding model’) to the query. A combined prompt is then created from the query and the retrieved information, improving the output from the generative network. f) RAG with context compression using xRAG (*33*). A small network, trained with the LLM, maps vectors from the text embedding model into a single token representation (a token is the atomic unit an LLM operates over, i.e. a ‘chunk’ of characters), which is fed into the first layer of the LLM together with the tokens of the query.

We suggest that hippocampal-neocortical interaction resembles RAG, but crucially with additional compression and consolidation mechanisms. Memories in the hippocampus are stored in a compact conceptual form, as in the LLM literature on context compression (e.g. (*33*)). Compression is a natural consequence of schema learning; schemas develop in neocortex over time, and thus feature in hippocampal traces which bind together neocortical representations. Consolidation then gradually trains the neocortical generative network on memories from the hippocampus. The conceptual representations in hippocampus result from compressing experience in a way that is optimal for efficient reconstruction of memories by neocortex, and all recall of hippocampal traces is retrieval-augmented generation, requiring the generative network to fill in the outline of a memory with its predictions.

The ‘context’ (or ‘prompt’) provided to an LLM can be thought of as its ‘working memory’ (*34*), with the limited context length of an LLM representing the limited capacity of biological working memory. Note that the neocortical model’s outputs could be prompted by perception rather than memory using the same mechanism.

### Previous work

There is a long history of examining how short, medium, and long-term stores support encoding, consolidation, and recall across the ‘lifespan’ of a memory (e.g. (*42*)). Crucially, semantic memory is thought to be extracted from episodic memory through consolidation (*2, 3, 43*), learning statistical relationships from experience to aid prediction and thus survival (*44*). But episodic memories also feature conceptual representations from semantic memory (*45, 46*); to use the terminology of Piaget (*47*), the encoding of memories using existing schemas reflects ‘assimilation’, while the updating of schemas from memories reflects ‘accommodation’. Both semantic and episodic memory can interact within working memory when reconstructing what happened in the past or imagining what will happen in the future (*48*), with episodic memory contributing specifics and semantic memory contributing generalisable knowledge (*49*). The conception of memory as (re)constructed rather than veridical (*20*) highlights the potential of recently developed deep generative networks to shed new light on these questions.

The idea that memories are rapidly encoded by the hippocampus and then replayed over the course of systems consolidation to train a generative or predictive model of the world in neocortex has received much recent interest. The combined system is thought to support multiple cognitive functions including episodic memory, semantic memory, imagination, and inference (*27, 30, 38, 50, 51*). This provides a mechanistic account of the theory that episodic memories are reconstructions that are influenced by our beliefs, i.e. that recall involves ‘predicting’ the past (*37, 52, 53*). However previous work has focused on static patterns, whereas episodes are sequential, and arguably the most crucial function of semantic knowledge involves prediction of what will come next.

Memory helps to predict the future, not just recall the past (*44*). Planning often requires anticipating the outcome of a possible action, e.g. the sequence of future states and rewards that would follow it. The mental simulation of future states to aid decision-making is known as ‘model-based’ planning (*54,55*), which improves with consolidation (*56*). We suggest that the generative network trained on memories can simulate events to support behaviour, such as when deciding between possible actions, or when inferring the next state given the sequence so far. (See also (*57*).)

## 2 Results

In Section 2.1 we explore the compression, encoding, recall, consolidation, and forgetting of narrative events. We show how each event can be efficiently encoded in the hippocampus as a conceptual code plus surprising details, from which the neocortex reconstructs the event during recall via ‘retrieval-augmented generation’, with these reconstructions gradually consolidated into neocortex, where interference from new consolidation produces forgetting. We show from simulations how this produces the changes observed in human memory over time.

In Section 2.2 we explore how the model captures the influence of semantic memory on episodic memory, and show that characteristic distortions towards the ‘priors’ of the network arise as a consequence, which captures gist-based distortions seen in human memory. In Section 2.3 we show that consolidation of memories into the neocortical network enables learning of shared structure, which can be used to solve new problems. We simulate human behaviour in two non-linguistic relational inference tasks, involving the prediction of new spatial or family relationships. We also show that human-like problem solving is supported by the mapping of new tasks to a learned representation in the hidden layers.

We model the hippocampal network as a sequential variant of the modern Hopfield network (MHN) for recalling state 𝑛 + 1 given state 𝑛 (*31, 58*), coupled with a MHN for retrieving state 𝑛 from a noisy or partial version. Given a state as a cue, recall is simulated by finding the nearest stored state with the latter MHN, and retrieving the rest of the sequence with the former. For further details see Section 4.2.2. Replay is simulated by randomly sampling from the set of memory units corresponding to the first state of the sequence, and retrieving the rest of it. Sequences are repeated many times during training, reflecting offline replay rather than online learning from a single exposure. (Note that after confirming that the model hippocampus can recapitulate sequences in each task, we avoid unnecessary computational cost by using these sequences directly, unless stated otherwise below.)

### 2.1 Modelling hippocampal-neocortical interaction as retrieval-augmented generation

We begin by simulating the compression, encoding, and consolidation of memories, and the differing mechanisms for recall over the course of a memory’s lifespan.

#### 2.1.1 The compression and encoding of episodic memory

In our model memories are compressed for efficient storage (*59, 60*). Neocortex captures the gist of the episode, which is stored in hippocampus together with unpredictable details. The stored hippocampal gist vector and additional details are unpacked by neocortex into a full description of the episode during recall. This explains not only the presence of conceptual representations in hippocampus (*45*), but also the fact that memories are schematised from the time of encoding (*61, 62*).

Retrieval-augmented generation is used to reconstruct memories or solve new problems based on this compressed information, with hippocampal traces retrieved based on relevance to the current task or prompt, and loaded into the context of the generative network. This minimises the hippocampal storage of redundant information that is already well-predicted by the generative network (see also (*27*)). In other words, this hippocampal representation allows the neocortical network to (re)generate experiences in a more efficient way, and the conceptual representations are optimised to enable generation. We simulate this by following recent LLM research on ‘context compression’, aimed at mitigating the practical problems of dealing with very long prompts by transforming them into vectors which capture their meanings (*33, 63*).

Here we use the Mistral-7B-Instruct-v0.2 (*29*) LLM to represent the neocortical network, together with pre-trained weights from (*33*) to transform episodes into compressed xRAG vectors. The small compression network loosely represents the neocortex’s ability to perform dimensionality reduction (*64*) and produce gist-like representations of events (*65*). This is one of several approaches to context compression which learn a compact vector representation of text (see also (*63*)), and does not require any special training of the LLM itself - just additional weights that map the vector representation of a sequence (i.e. the output of a ‘text embedding’ model) to a token in the input layer of the LLM (see Figure 1f). Note that tokens are chunks of co-occurring characters that make up the basic units processed by an LLM.

To illustrate how gist, detail, and semantic memory interact, we simulate the encoding and recall of narrative events (Figure 2), using 500 stories from the ROC Stories dataset (*66*). To encode a story in compressed form, its embedding is obtained from the text embedding model (*67*), from which the xRAG vector is derived via the xRAG weights. Each story is encoded as its xRAG vector plus unpredictable details in the model hippocampus. To identify unpredictable details, all phrases are extracted from the text, then the perplexity (a measure of prediction error) of each phrase given the xRAG vector is calculated. In other words, for each phrase, the model is prompted with the xRAG vector and the phrase in question. Then the phrases are ranked in descending order of perplexity, with the top *n* phrases stored as the unpredictable details. See Section 4.2.1 for further details.

**Figure 2:**
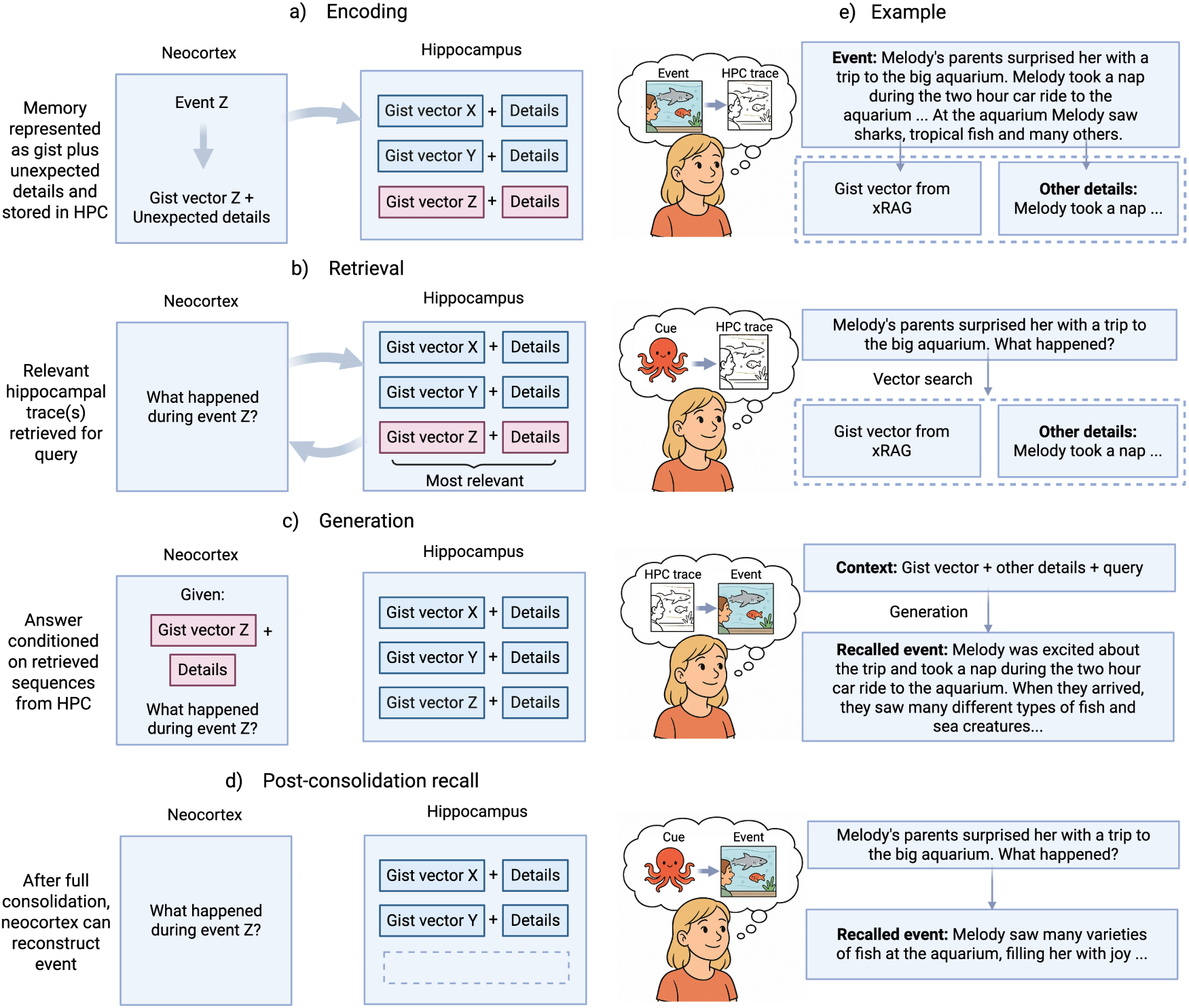
A model of hippocampal-neocortical interactions as retrieval-augmented generation with stories. a) The neocortex generates a gist vector for the narrative, which is encoded in the hippocampus together with unpredictable details. b) Given a query as the input to recall, the neocortex searches the hippocampus for relevant traces, e.g. by finding nearby gist vectors. c) The generative network in neocortex produces an ‘answer’ conditioned on the retrieved hippocampal trace(s). d) Once a story is fully consolidated, the hippocampal trace is no longer needed to approximately reconstruct the event. e) The stages of encoding and recall are shown for an example story from the ROC Stories dataset (*66*), with Mistral-7B-Instruct-v0.2 used to simulate the neocortex, and xRAG vectors (*33*) representing the conceptual gist.

Given a query as the input to recall, the neocortex ‘searches’ the hippocampus for relevant traces. This is simulated by producing an xRAG vector for the query (as described above), and then by retrieving the trace(s) with the nearest xRAG vector(s) from the model hippocampus (see Figure S1). The context given to the LLM consists of the xRAG vector(s), the details, and a prompt describing the task. Note that the xRAG vector only occupies the space of a single token in this input. The final recalled story is the output of the LLM.

Figure 2e and Table S1 show examples of encoded and recalled stories. For instance, a story about a girl’s visit to an aquarium is encoded as its gist vector and a surprising detail (that the girl took a nap on the journey). The model neocortex reconstructs an approximation of the original story from the gist vector and the detail, with a few semantic intrusions added to the story, e.g. ‘Melody’s favorite part was the shark tank’; these intrusions are shaped by the expectations of the model neocortex, as in (*20*). This hints at how gist, detail, and semantic memory can interact in the model during recall.

The encoding of gist plus details was also compared to three baselines: encoding the story in full detail, encoding only the gist, and simply imagining the continuation (i.e. without using the model hippocampus at all). The mean size per memory (in tokens) and the accuracy of recall (the cosine similarity between the embeddings of the original and recalled story) were compared across these conditions, showing a trade-off between efficiency and accuracy in memory in Figure 3a. The gist plus details encoding strikes a balance between these objectives.

**Figure 3:**
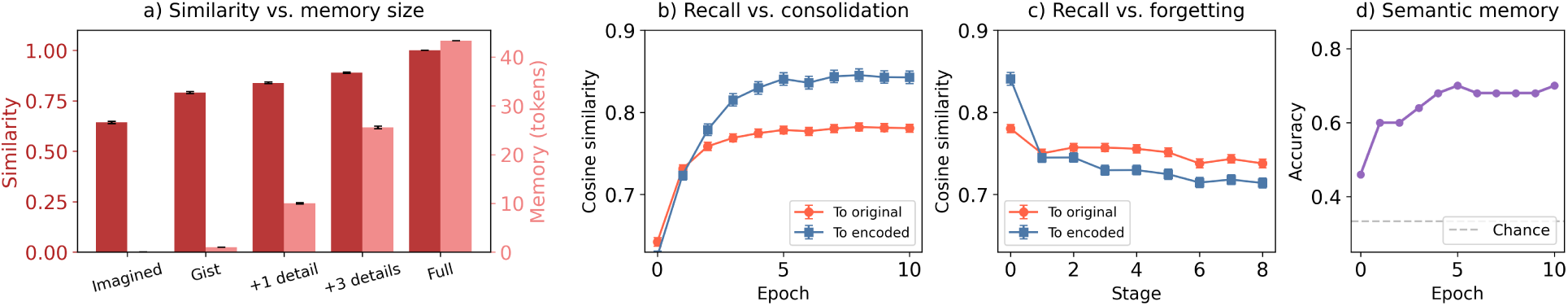
Encoding, consolidation and forgetting of narratives (ROC stories (*66*)). a) The size of the memory trace stored in the model hippocampus (in tokens) and recall accuracy (cosine similarity between original and recalled stories’ embeddings with ‘all-MiniLM-L6-v2’; (*69*)) for five variants. From left to right: simply imagining the event based on background knowledge, storing only the gist vector, storing the gist vector with one surprising detail, storing the gist vector with three surprising details, and storing the event in full. b-d) How neocortical representations change over time. b) The cosine similarity in embedding space between the neocortical memory and the original story (red), and between the neocortical memory and the replayed version (blue), which is itself a reconstruction of the compressed encoding. As the story is consolidated, the neocortex’s reconstruction moves closer to the encoded story in embedding space. c) As new memories are consolidated into neocortex, forgetting occurs (we assume the hippocampal trace has faded away). Over the course of new memory consolidation (50 new memories per stage), the neocortex’s reconstruction moves away from the encoded story in embedding space. d) Semantic memory accuracy (fraction of 100 questions for which the model’s answer is most similar to the correct answer) before and after consolidation.

#### 2.1.2 Modelling consolidation and forgetting

In our model sequential memories are initially encoded in the hippocampus in compressed form, reconstructed by the neocortex through ‘retrieval-augmented generation’ (*32*), and then gradually consolidated into a generative model in neocortex, through prediction error minimisation on the replayed memories. This provides a mechanism for the training of neural ‘foundation models’ (*24*), which can both capture specific memories *and* learn more general schematic knowledge to enable problem solving.

500 stories from the ROC Stories dataset (*66*) were encoded in the model hippocampus as described above (here in gist form without additional details for greater interpretability). To simulate consolidation, the memories were reconstructed from these traces, and the generative network was trained on the result (using low-rank adaptation; (*68*)). As the story is consolidated, the neocortex’s reconstruction moves closer to the encoded story in embedding space (Figure 3b), but distortions in the reconstruction are still observed. (See Section 2.2 for a more extensive exploration of distortions in our model.)

We also tested semantic memory for the content of the narratives by asking factual questions about 100 stories from the dataset, e.g. ‘Remember the story in which ‘Jane had recently gotten a new job.’ Why did Jane arrive late on her first day?’. Each prompt contains enough context to pick out a single relevant story, and the question has an answer in that story. Semantic memory is quantified as the fraction of questions for which the output is more similar to the correct answer than two alternatives (based on cosine similarity in embedding space). Performance increases with consolidation, showing that the network does not simply overfit to the exact text but can abstract semantic information from the story (Figure 3d and Table 3).

To simulate forgetting in the neocortical network, we tested the recall of the consolidated memories as the network was trained on new stories (representing the consolidation of new memories, in the absence of hippocampal reminders of the old memories). Over successive stages of new memory consolidation, the neocortex’s reconstruction moves away from the encoded story in embedding space (Figure 3c). See Table 2 for an example of the same event’s neocortical reconstruction accuracy improving during consolidation and declining during forgetting.

**Table 2:**
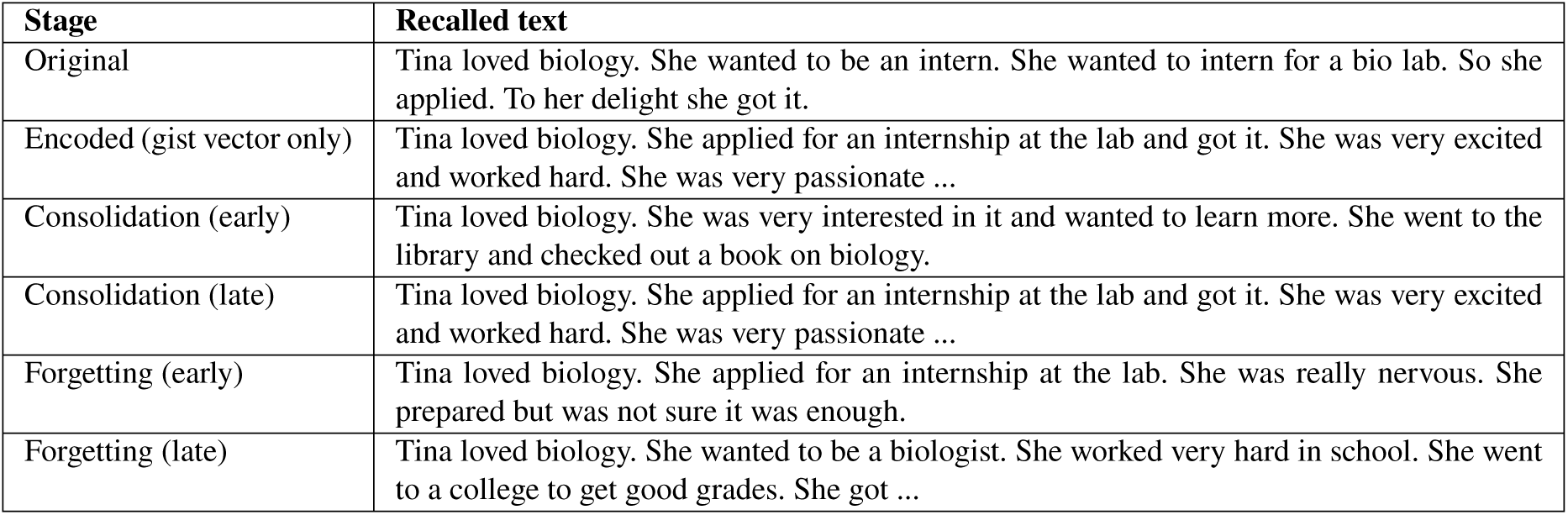
Examples of gist-based encoding, consolidation and forgetting. Text for a randomly selected story across six stages: the original story, the memory as reconstructed from the compressed hippocampal trace, the neocortical memory early and late in consolidation, and the neocortical memory early and late in forgetting. The initially encoded hippocampal memory produces gist-based distortions, and the neocortex gradually learns this event through consolidation, further distorting it. The neocortical memory gradually fades as newly encoded memories interfere. Note that when the story is initially encoded, only the gist (xRAG) vector is stored in the model hippocampus in this simulation, which increases the bias towards prior experience, but in the full model surprising details would be encoded too.

| Stage | Recalled text |
| --- | --- |
| Original | Tina loved biology. She wanted to be an intern. She wanted to intern for a bio lab. So she applied. To her delight she got it. |
| Encoded (gist vector only) | Tina loved biology. She applied for an internship at the lab and got it. She was very excited and worked hard. She was very passionate ... |
| Consolidation (early) | Tina loved biology. She was very interested in it and wanted to learn more. She went to the library and checked out a book on biology. |
| Consolidation (late) | Tina loved biology. She applied for an internship at the lab and got it. She was very excited and worked hard. She was very passionate ... |
| Forgetting (early) | Tina loved biology. She applied for an internship at the lab. She was really nervous. She prepared but was not sure it was enough. |
| Forgetting (late) | Tina loved biology. She wanted to be a biologist. She worked very hard in school. She went to a college to get good grades. She got ... |

**Table 3:**
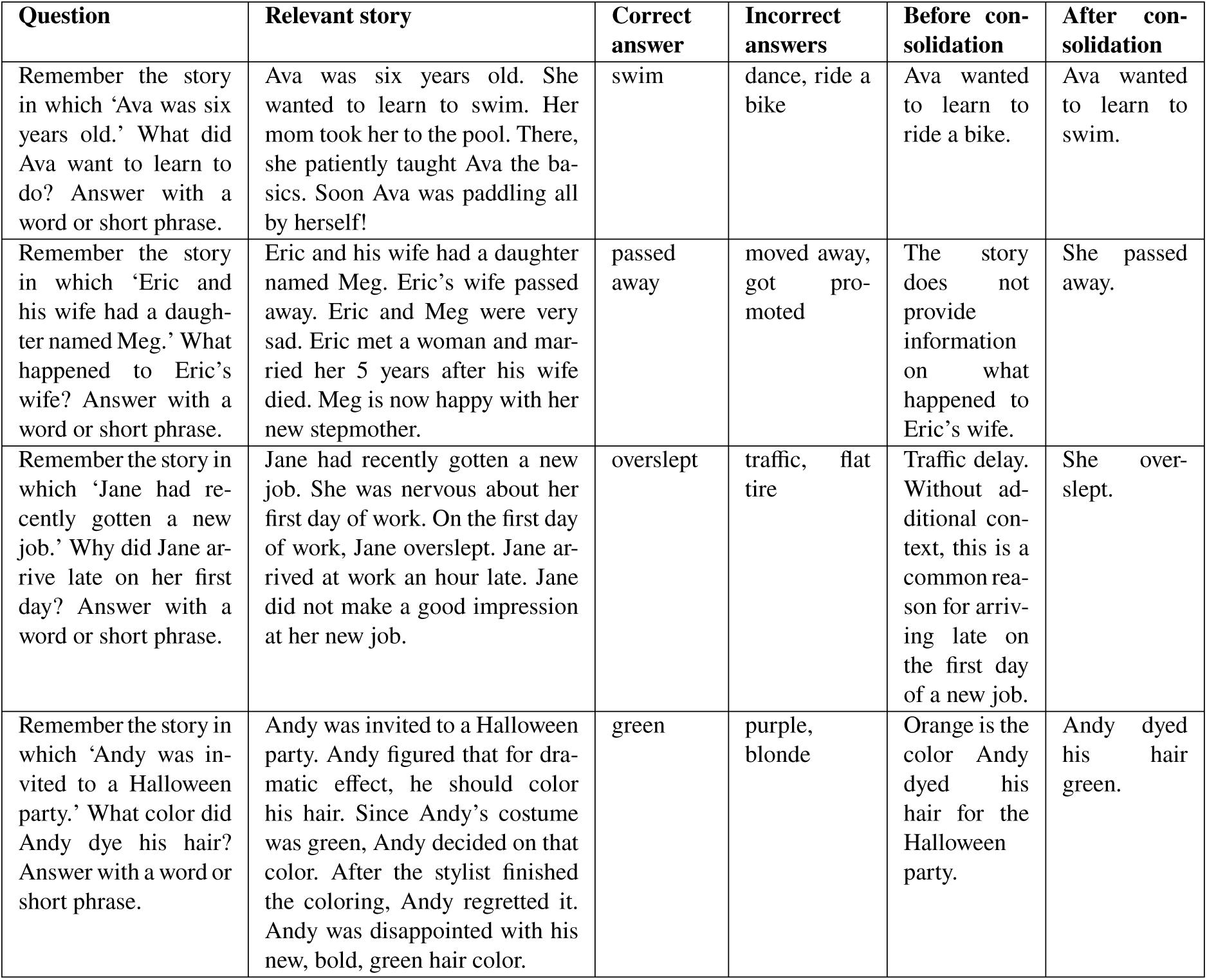
Neocortical semantic memory for particular events is acquired through consolidation. Neocortex-only answers for selected semantic memory questions before and after ten epochs of consolidation of 500 ROC stories (*66*). The model neocortex is prompted with the ‘Question’. Its answer is marked as correct if it is more similar to the ‘Correct answer’, as stated in the ‘Relevant story’, than the ‘Incorrect answers’ (as measured by the similarity of their text embeddings). Note that in the full model, semantic memory would rely on RAG before consolidation, with the relevant event retrieved into the context from hippocampus, but here only the neocortical memory is shown.

Putting these results together suggests an account of the inaccuracy of human episodic memory. To start with, the encoded version of an event is already highly compressed, relying on schemas from the neocortex. Then as consolidation progresses, memory becomes less dependent on the hippocampal trace, and more dependent on the neocortical reconstruction. We assume that the hippocampal trace is no longer maintained once consolidated into neocortex, with consolidation continuing until the trace can be *approximately* recon-structed by neocortex, rather than until it can be *perfectly* reconstructed. The consolidated version is therefore even less accurate than the encoded version. In addition, once fully consolidated into the neocortical model forgetting occurs due to interference.

#### 2.1.3 Changes in memory content over time

The HIPPOCORPUS dataset (*18*) contains participants’ written descriptions of recalled autobiographical events, with a subset of the recalled events retold months later. We analysed it to track how concrete vs. abstract, how rich vs. poor in detail, and how specific vs. general memories were for more recent and more remote episodic memories. We then compared the original, encoded, and consolidated versions of stories in our simulations using the same metrics. This was achieved by using an LLM as an annotator (gpt-4o-mini via the OpenAI API).

Figure 4d shows that the more remote memories are less concrete, rich in detail, and specific, in line with many similar findings. For comparison, we simulated the process of encoding and consolidation in our model with the ROC Stories dataset (*66*) as described above, because the HIPPOCORPUS dataset does not contain ‘ground truth’ descriptions of the remembered events. Figure 4a shows that the encoded memories are significantly less concrete, detailed, and specific than the original stories, and the consolidated stories score even lower on these attributes. In Tables S3 and S4, SI, we show that reduction in these attributes does not simply reflect less memory being recalled by controlling for length, confirming that there are still significant differences. In addition, Figures 4e and g show that more commonly used words occur in the remembered stories. Furthermore, memory concreteness, richness, and specificity decline as forgetting proceeds (Figure 4b).

**Figure 4:**
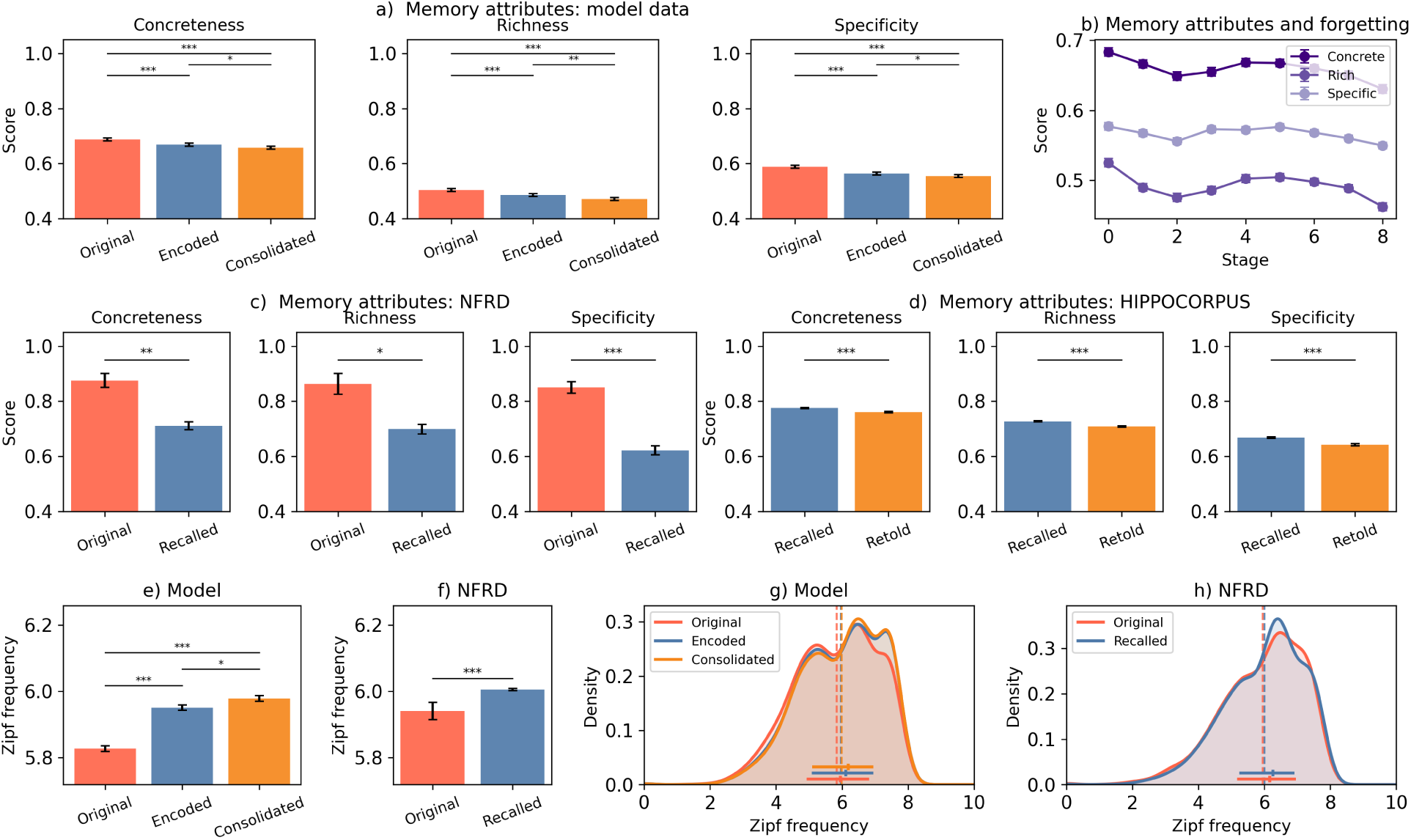
How memory content changes over time. a) Memory concreteness, richness in detail, and specificity (as scored by gpt-4o-mini) for the original ROC Stories data (*66*), the memories based on the compressed hippocampal encoding, and the memories reconstructed by neocortex alone after consolidation. b) Memory concreteness, richness, and specificity decline as forgetting proceeds. c) As in a) but for original and immediately recalled stories in the Naturalistic Free Recall Dataset (NFRD; (*19*)). d) As in a) but experimental data for ‘recalled’ stories, and the same stories ‘retold’ at a later date from the HIPPOCORPUS dataset (*18*). e) The mean word frequency (across English language text, computed with the ‘wordfreq’ Python library) for words in the original, pre-consolidation, and post-consolidation stories. f) Likewise but experimental data for the NFRD texts. g) The distributions of word frequencies for the categories in e), with means shown as dotted lines and medians and interquartile ranges shown as bars, showing increased use of high frequency words after encoding and consolidation. h) Likewise but experimental data for the NFRD texts.

The Naturalistic Free Recall Dataset (NFRD; (*19*)) consists of transcripts for four stories recalled by more than one hundred participants each, immediately after listening to the story. We repeated the analysis above for the original and recalled stories for comparison to the original and encoded versions in our simulation, observing a similar decrease in those attributes (Figure 4c), and furthermore the use of more common words in the recalled NFRD stories (Figure 4f and h). (We note that schematisation is not the only explanation for vocabulary changes; even with a perfect memory for an event, one might paraphrase its content in more familiar language.)

### 2.2 Modelling the interplay of semantic and episodic memory in consolidation

Section 2.1 sketched out how narrative memories change over time, but now we explore the effect of existing schemas on memory in more depth. Here we investigate how the model distorts memories towards expectations, consistent with human data.

#### 2.2.1 Gist-based distortions and prior experience

In the Bartlett experiment (*20*), students heard a story called ‘The War of the Ghosts’ and were asked to recall it after different time intervals. The recalled story was distorted to be more consistent with the students’ background knowledge of the world, with distortion increasing over time (*70*) (Figure 5b). To simulate encoding, the story was compressed as described above (here with no poorly predicted details for greater interpretability). To simulate consolidation, Mistral 7B was then trained on its own reconstruction of the encoded story. Recall of the story was explored by giving the network its first few words (‘One night two young men from Egulac’), and inspecting the predicted continuation. The consolidated story displayed even more distortion than the encoded version (Figure 5e), suggesting that consolidation of a compressed memory is an ‘approximation of an approximation’.

**Figure 5:**
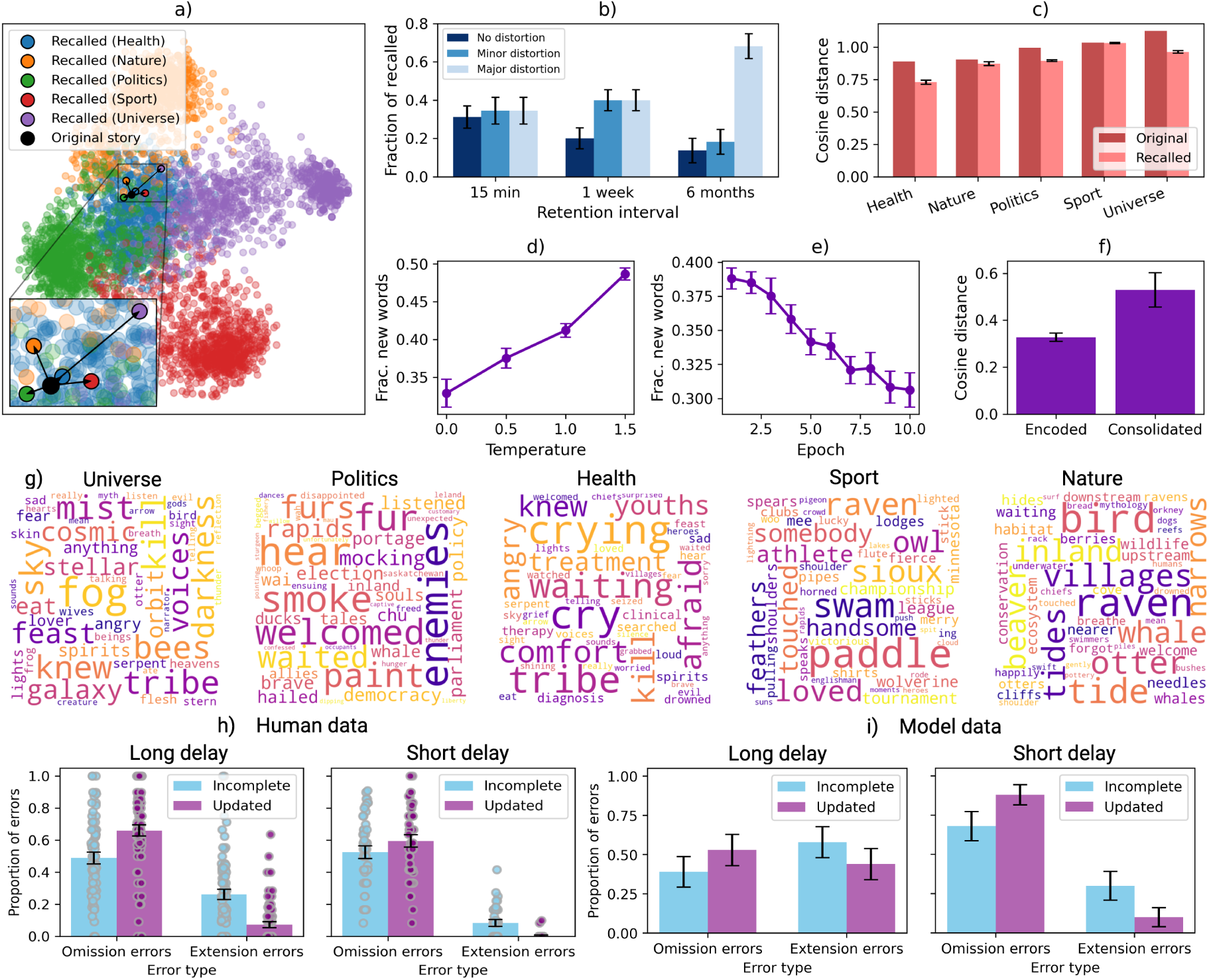
The effect of prior knowledge on narrative distortions. a) The neocortical network (pre-trained Mistral 7B) is fine-tuned on Wikipedia content (*71*) from each of five categories plus the Bartlett (*20*) story, (with the original rather than compressed version ‘replayed’ to isolate the effect of consolidation in parts a, c-e, and g). The embeddings of the training data plus the recalled stories for each model (at a temperature of 0.5) are obtained, using ‘all-MiniLM-L6-v2’ (*69*). These embeddings are projected into 2D with PCA, showing that recall moves each story towards that network’s training data. b) Data reproduced from the Bergman and Roediger (*70*) replication of Bartlett (*20*) showing that the fraction of recalled propositions in the story with major distortions (defined as showing semantic influences) increases at greater delays in a repeated reproduction experiment. c) The cosine distance (1 - cosine similarity) between the mean embedding for each category and the original story (purple) and recalled stories (orange). Recalled stories become more similar to the background dataset. Error bars give the SEM for the 200 recalled stories sampled. d) The effect of temperature on the number of new words in the recalled Bartlett stories (‘semantic intrusions’). Error bars give the SEM. Results are for the model at epoch five. e) The effect of epoch on the number of new words in the recalled Bartlett stories, where an epoch is one presentation of the entire training dataset. Error bars give the SEM. Results use a temperature of 0.5. f) The mean cosine distance between the recalled and original stories before and after consolidation (at a temperature of 0.5, and with ten epochs of consolidation). g) Word clouds showing ‘semantic intrusions’ (produced with the ‘wordcloud’ Python package) at a temperature of 0.5, aggregated across five trials. h) Event extension and contraction results, showing omission errors and extension errors after long (left) and short (right) retention intervals, adapted from Raykov et al. (*21*) with permission. i) The proportion of stories (200 per type) recalled with omission and extension errors, both before consolidation (cf. short delay in h) and after (cf. long delay). As in part h), error bars give the 95% confidence intervals of the mean.

To explore the effect of the model’s ‘priors’ (i.e. its expectations based on previous learning) on recall of narratives, the Bartlett story (*20*) was consolidated after pre-training on ‘background data’. A dataset of Wikipedia content (*71*) was used, with five categories of article selected (‘Politics’, ‘Health’, ‘Universe’, ‘Sport’, and ‘Nature’). The training data for each model was made up of 1000 articles sampled from the relevant category (with the first 1000 characters of each article taken). The model was trained for 10 epochs on this dataset, where each epoch is one full presentation of the training set. Next, the Bartlett story was consolidated into the network in the same way (with the original story used for training to isolate the impact of consolidation rather than encoding). The temperature for sampling continuations from the model was also varied between 0 and 1.5 (see Section S1.1.2).

When the Bartlett story is ‘consolidated’ into the generative network memory distortions are observed, as in the human data (see Table 4). Distortions in recalled stories reflect the ‘priors’ of the generative network. The word clouds in Figure 5g show that new words added to the story (i.e. ‘semantic intrusions’) are representative of the background dataset used, and Figure 5a and c show that the recalled stories move closer towards the background dataset in text embedding space. See also Table 4 for selected examples of semantic intrusions from the five models. Whilst distortion occurs even with ‘greedy decoding’, more ‘semantic intrusions’ are observed at higher temperatures, as Figure 5d and Table S5 show.

**Table 4:** Examples of the effects of prior knowledge on narrative distortion. a) Extracts from retrievals of the Bartlett (*20*) story for networks trained on different categories of prior knowledge (temperature = 1.0), showing how semantic intrusions reflect the priors of the generative network. b) Event extension and contraction examples before consolidation showing the effect of encoding with compression. The selected incomplete and updated stories (which are of the same length) display extension and omission errors respectively. The model is prompted with the first 150 characters of each story.

| a) Semantic intrusions in consolidated memory reflect priors |  |  |
| --- | --- | --- |
| Nature | The intruders lit a fire and started eating berries. |  |
|  | a great herd of caribou . . . crossed the river |  |
|  | The Orcas were hunting humans that night. |  |
| Health | The young men ran home and told this to their people and the doctor was called |  |
|  | While they were crossing the river in their canoe, it overturned and both men were drowned. |  |
|  | The spirits gave the two youths gifts, including a powerful medicine pipe |  |
| Sport | [He] took up his bow and arrow, for he was the greatest hunter of the world |  |
|  | Men selected, not only for their bravery, but for their swimming ability . . . |  |
|  | They were good hunters. |  |
| Politics | [They] decided to join their forces and fight against the common enemy |  |
|  | the man who was in charge . . . addressed them in their own dialect |  |
|  | —The boy’s father, who was a chief, sent his people away to hide |  |
| Universe | He told them how the moon came to be and how the stars got their positions in the sky. |  |
|  | It told of how the world was once covered in water, then the land appeared |  |
|  | They saw in a dream a vast multitude of stars falling from the sky. |  |
| b) Event extension and contraction examples before consolidation |  |  |
| Type | Original | Recalled |
| Typical | Stephen wanted a cardigan for his birthday. Sally bought one for him. She wrapped it up. She wrote a note. She gave it to him! The end. | Stephen wanted a cardigan for his birthday. Sally bought one for him. She wrapped it up. She wrote a note and gave it to him! The end. |
| Incomplete | Floyd was enjoying a summer day of inspecting cicada shells. He found many of them and hung them on his shirt like decorations. He squatted at a large tree’s roots when he spotted a really nice one. When he picked it up, t The end. | Floyd was enjoying a summer day of inspecting cicada shells. He found many of them and hung them on his shirt like decorations. He squatted at a large tree’s roots when he spotted a really nice cicada. He decided to add it to his collection. The end. |
| Updated | Kris lost her job. She was sad at first. She looked for a new job. She got an interview. Kris got a new job! Jack had always wanted his own dinosaur. He asked Santa Claus for one every year. He never would get one and beca The end. | Kris lost her job. She was sad at first. She looked for a new job. After an interview Kris got a new job! The end. |

#### 2.2.2 Event extension and contraction

Distortions can apply to the structure as well as the content of narratives. Event extension is the tendency to extend certain events in memory, while event contraction is the tendency to curtail them. Raykov et al. (*21*) demonstrate this with three types of video: complete videos end at a natural event boundary, incomplete videos are curtailed before this point, and updated videos are extended beyond it. They find that incomplete videos are often extended in memory (extension errors), while updated videos are often shortened (omission errors), and show that this is observed at long and short delays (Figure 5h).

We simulate the effect of consolidation in (*21*), using simple stories in text form (*72*). The majority are ‘typical’ stories, unmodified from the ROC Stories (*72*) dataset, whereas an ‘incomplete’ story has characters removed, and an ‘updated’ story has characters added, so it continues beyond its natural ending. The data consisted of 200 typical stories, 200 incomplete stories, and 200 updated stories, where ‘updated’ and ‘incomplete’ stories are all the same length. First the stories were encoded in compressed form as described above, and pre-consolidation recall was tested by giving the first 150 characters of each story as a query for retrieval-augmented generation. Then the *reconstructions* of these compressed memories were replayed to simulate consolidation; Mistral 7B (*29*) was fine-tuned on the reconstructed stories for three epochs, and its recall tested. Both before and after consolidation, omission errors are more common in updated than incomplete stories, while extension errors are more common in incomplete than updated ones (Figure 5i). By bringing the story to a more natural conclusion or by removing an incongruous ending, the stories are made more similar to the network’s expectations. See Table 4 for examples. This mirrors findings on boundary extension and contraction in memory for images (*73, 74*); in both cases, memories are distorted towards a prior encoded in the network by its previous ‘experience’, consistent with Bayesian views of memory (*27,52*).

### 2.3 Neocortical learning of sequential structure for problem solving

Next we investigate how consolidation develops generalisable knowledge which supports problem solving. To enable simulations of training from scratch, we use a smaller LLM architecture, GPT-2 Medium (*23*), and simplify the model by storing memories in full rather than compressed form.

The ability to anticipate future observations improves with consolidation (*75–77*), leading to downstream improvements on a diverse range of tasks even as perceptual details fade. We demonstrate how the neocortical network can extract sequential structure from memories through consolidation using the example of structural inference, which also the model’s application to non-linguistic stimuli. A spatial example of structural inference is the finding of shortcuts, as this relies on the common structure of space, and a non-spatial example is inferring that A is the grandfather of C from the knowledge that A is the father of B, and B is the father of C, which relies on the common structure of family trees. The relations in these tasks can be seen as edges in graphs, so that multiple tasks with a common transition structure can be simulated using a fixed graph with different nodes for each task (*22*).

We simulate how consolidation enables structural inference by the neocortical network. We consider inference in two types of graph: a spatial graph and a simple family tree graph (as in (*22*)). We trained GPT-2 models on random walks on these graphs, corresponding to sequences of observations encoded in the hippocampus, and then tested the models’ inference abilities on novel sequences with the same underlying structure.

In the spatial graph, a three-by-three grid represents a simple 2D environment, where the nine nodes are locations and the edges between them (‘NORTH’, ‘EAST’, ‘SOUTH’ and ‘WEST’) are possible movements (Figure 6a). Whilst each graph’s structure is the same, nodes are labelled with names (random pairs of letters) to represent arbitrary features at a particular location. Trajectories through the environment are walks on the resulting directed graph, which are represented as strings such as ‘ab EAST wd SOUTH ea WEST hn’. (See graph transformers for an alternative approach (*78*).)

**Figure 6:**
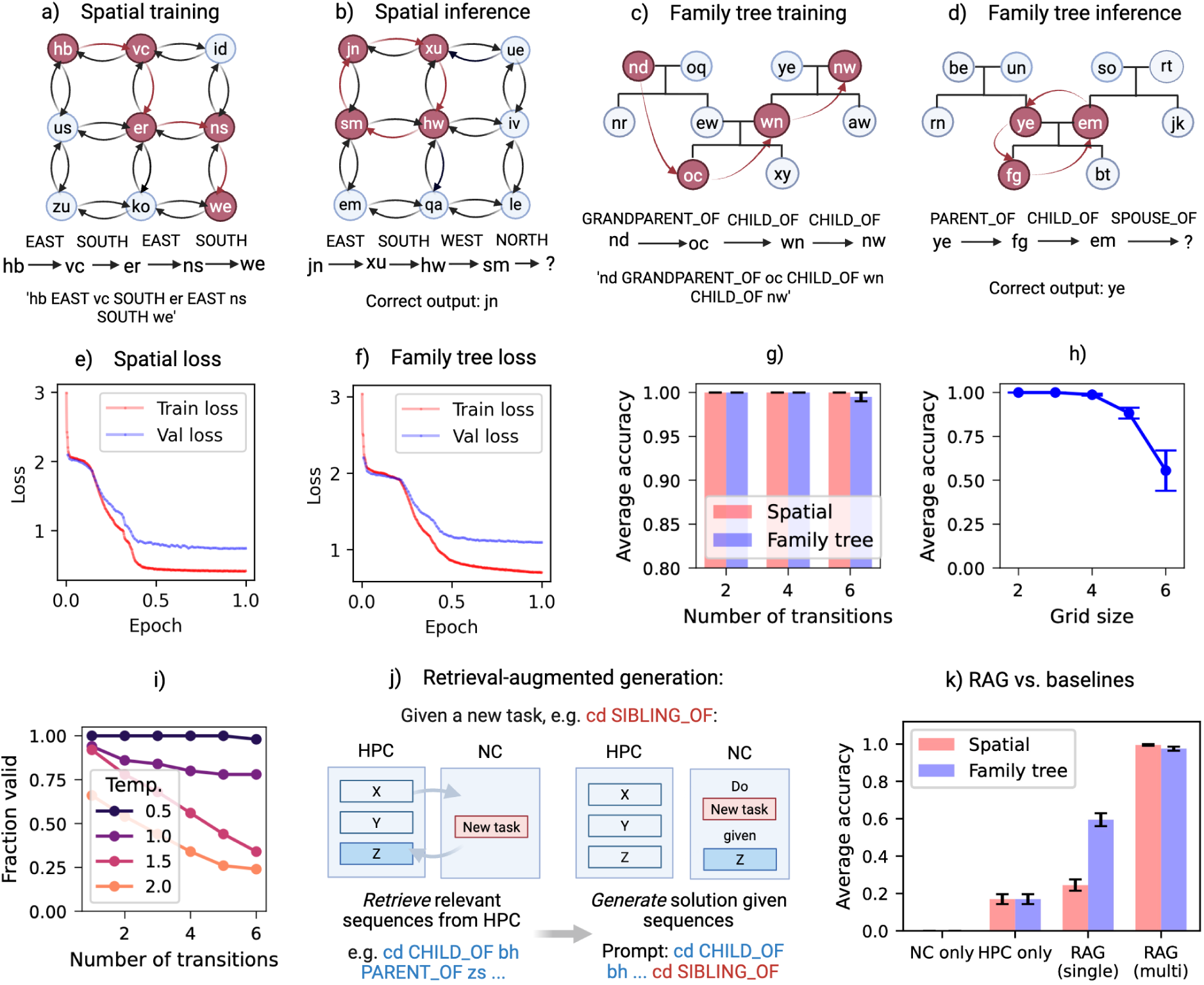
Learning structural regularities. a-b) The spatial graph task. a) Spatial trajectories are modelled as walks on the graph, and represented as strings. b) Structural inference is tested as the ability to complete sequences on unseen graphs based on structural regularities, e.g. inferring that the next location in the sequence shown is ‘jn’. c-d) As above but for the family tree graph task. e) The training and validation losses (i.e. aggregated prediction errors) of GPT-2 trained for one epoch on sequences from 100,000 spatial graphs. f) As above but for the family tree graph task. g) Inference, as shown in parts b) and d), was tested using sequences from *novel* environments / family trees in which the final entity can be inferred from the sequence so far. Performance is grouped by the number of transitions. Error bars give the SEM. h) Generalisation to different grid sizes. The spatial inference model - trained on paths in a 3×3 grid of locations - was tested on a range of square grid sizes (from 2×2 to 6×6), and generalises beyond its training data. i) The fraction of ‘imagined’ spatial trajectories (as generated by the spatial inference model when given a random location as an input) which are structurally valid, at a range of temperatures. A sequence is marked as invalid if different locations are found at the same inferred co-ordinates (e.g. ‘sz WEST eq EAST zr’). j) RAG applied to an inference task. Relevant sequences are retrieved from the hippocampus, and then provided as input to the generative network together with the new task. k) Inference accuracy for the spatial and family tree inference tasks with RAG, in both a single-memory and multi-memory mode, and a hippocampus (HPC) only and neocortex (NC) only baseline for comparison.

The family tree graph has a simple structure for illustrative purposes, consisting of two children, their parents, and two sets of grandparents. See Figure 6c. We model this as a directed graph with edges ‘PARENT OF’, ‘CHILD OF’, ‘SPOUSE OF’, ‘SIBLING OF’, ‘GRANDPARENT OF’, and ‘GRANDCHILD OF’. As in the spatial case, all graphs have the same structure, but each graph has different names assigned to its nodes. Walks on the graph are represented by strings such as ‘lk PARENT OF nd SIBLING OF re’.

In each case, we created 100,000 graphs with the same structure but randomly chosen labels (pairs of letters) for the nodes. 20 random walks, of a number of transitions sampled from between 1 and 50, were gathered from each graph to create the training data, representing sequences of observations that might be experienced, encoded in the hippocampus, then replayed offline. GPT-2’s medium-sized architecture was then trained from scratch for one epoch (i.e. the full dataset was presented once). After training, we tested inference of the next node given the sequence so far, using greedy decoding of the output probabilities. For example, the next item after ‘uq NORTH sx EAST tp SOUTH ec WEST’ can be inferred to be ‘uq’, and the next item after ‘qk PARENT OF xm PARENT OF vw GRANDCHILD OF zq SPOUSE OF’ can be inferred to be ‘qk’. The sequences used for testing were sequences of relationships in *new* family trees or spatial grids.

Figure 6e and f show the ‘loss’ (aggregated error on the training data) of the spatial and family tree models respectively. In both cases the loss gradually decreases, indicating improved ability to predict the next node on the set of graphs used for training, which corresponds to the consolidation of previous experience. Tables S6 and S7 and Figure 6g show good performance on a range of *novel* structural inference tasks, involving sets of labels representing environments which have not been experienced before. Simpler inferences include inferring that going one step west then east takes one back to the original location, e.g. the correct continuation of ‘ab EAST cd WEST’ is ‘ab’. Similarly, a correct continuation of ‘ab PARENT OF cd CHILD OF’ is ‘ab’. But surprisingly complex inferences are also possible, e.g. that ‘kh CHILD OF oi SPOUSE OF tv CHILD OF fh SPOUSE OF xr GRANDPARENT OF gq SIBLING OF’ is followed by ‘kh’. Furthermore given a single new item the trained spatial model can ‘imagine’ new trajectories consistent with the learned graph structure (Figure 6i). We also tested inference on larger grids than the model was trained on to show that inference cannot solely be attributed to ‘rote memory’ of sequences of transitions (Figure 6h).

#### 2.3.1 Inference from memory as retrieval-augmented generation (RAG)

Episodic memories can be combined with general knowledge to support problem solving. Next we simulate how the generative network and hippocampal network could work together to achieve this via RAG.

In the simulations above, inference was tested as the continuation of a single new sequence in working memory. However working memory is limited (*16, 17*), so the brain must selectively recall relevant experiences. In addition, problem solving can synthesise information from more than one episodic memory retrieved from hippocampus; multiple fragments of information encoded at different times can be recombined with semantic knowledge, e.g. when inferring that A is the sibling of C from the observation on one day that A is the child of B, and on another that B is the parent of C.

We create a ‘toy example’ of retrieval-augmented generation using the two models in the structural inference results above, and consider a task in which multiple memories, from recent tasks which have not yet been consolidated, are required to infer the right answer (Figure 6j). For example, from the pair of memories ‘ab EAST cd SOUTH ef’ and ‘ef WEST gh’, it can be inferred that ‘gh NORTH ab’. 200 such tasks were created for each of the spatial and family tree experiments. To simulate RAG, the top two matches to the query (e.g. ‘gh NORTH ?’) are first retrieved from the model hippocampus. Then the model neocortex predicts the solution to the query with these retrieved sequences in context. We compare this to baselines of RAG with a single memory, the hippocampus only solution (randomly selecting one of the locations / people in the retrieved sequences), and the neocortex only solution (conditioning the generative network on the task alone, without retrieving sequences from the hippocampus). The results show that RAG supports structural inference based on hippocampal memories, whereas relying on either the hippocampal network or generative network alone gives worse results, and that the model can solve problems using information from multiple episodes (Figure 6k).

#### 2.3.2 Inspecting the hidden representations supporting inference

To understand how the neocortical network generalises to a new set of labels, we inspected representations in its hidden layers. Activations (vectors of length 1024) for each token of a random walk were extracted from the innermost layer (i.e. the 12th of the 24 transformer blocks) of the model neocortex trained in the spatial task above, and from the same layer of the GPT-2 medium model pre-trained on language and released by (*23*) as a baseline. Having obtained a vector for each location, we analysed the distances between these representations. We found that the nearer the points in the environment, the closer their hidden representations (Figure 7f). This is not true of the baseline model (Figure 7e). We then ran principal component analysis across representations extracted from random walks in many different environments. Figure 7b shows that our model has learned a shared structure across episodes, whereas this structure is not present in the baseline GPT-2 medium (Figure 7a); the same is true of the family tree model, in which the three ‘generations’ in the tree are separated in our model (Figure 7d) but not in the baseline model (Figure 7c).

**Figure 7:**
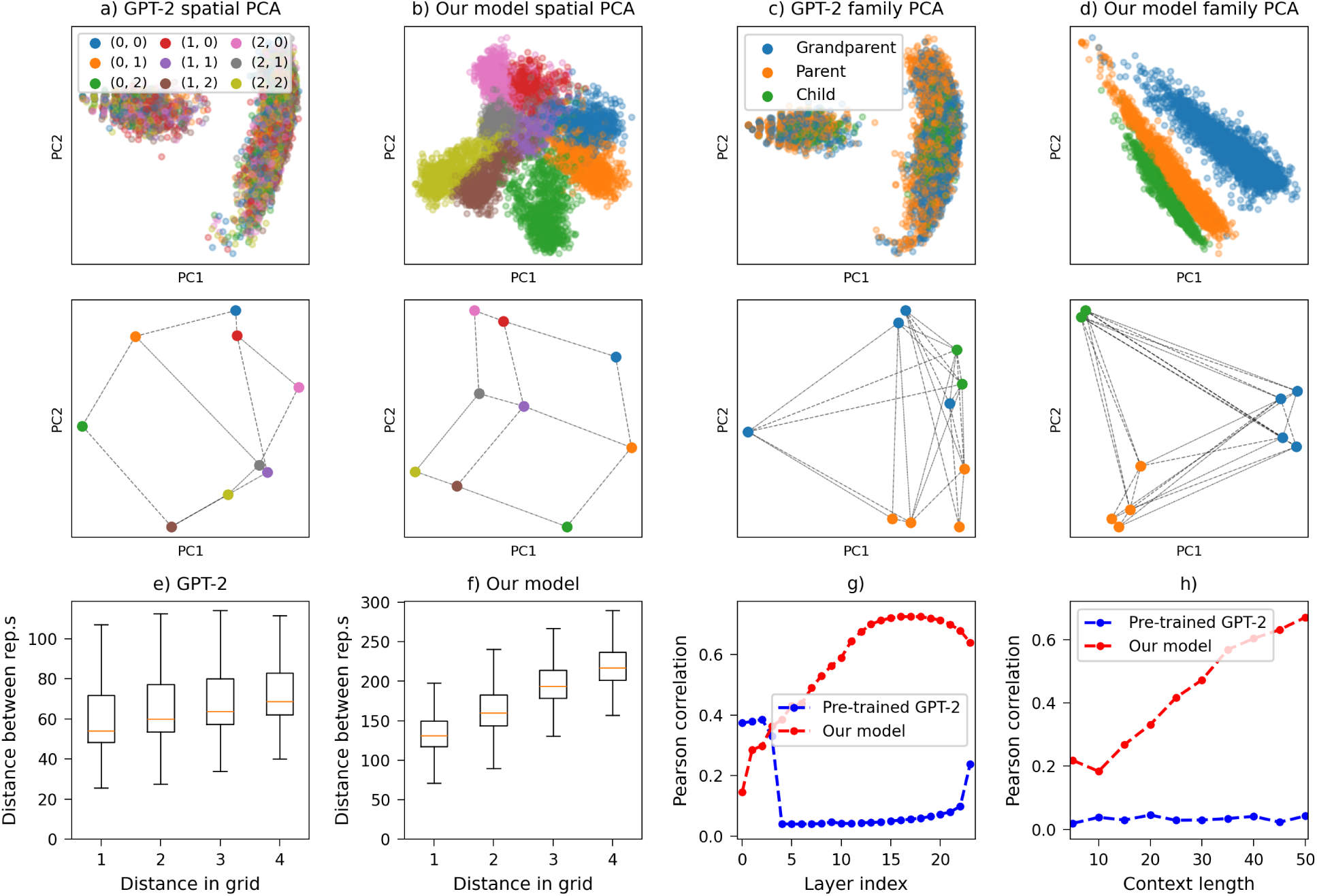
Inference is supported by learned neocortical hidden representations. a-b) Representations in the spatial inference model. Activations for each token of a random walk were extracted from the middle layer (12) to obtain a vector for each location. Principal component analysis was performed across representations extracted from random walks in many different environments (upper row), colouring each location by its co-ordinates in the environment, so that points with the same colour are different nodes which occupy the same position in the 3×3 grid (see example environments below, where dotted lines indicate possible transitions). GPT-2 medium pretrained on language is shown in a) and our model - GPT-2 medium trained on structural inference - in b). c-d) As above but in the family tree inference model. Here three colours indicate the three generations in the family tree. e) In the GPT-2 medium baseline, there is little relationship between Manhattan distance in the environment and Euclidean distance in token space. f) As in e) but for our model, showing that the token representations reflect distances in the environment. g) The Pearson correlation between Manhattan distance in the environment and Euclidean distance in each layer of our model (red) and GPT-2 medium (blue). h) The Pearson correlation between Manhattan distance in the environment and Euclidean distance in layer 12 for prompt lengths differing in number of transitions, for our model (red) and pre-trained GPT-2 medium (blue).

These results mirror the finding that a shared abstract representation of different spatial tasks with the same structure develops in cortex through consolidation, as has been shown using cross-map repetition suppression effects in fMRI (*79*). Once this abstraction of the task has developed, it supports reasoning based on information in the prompt, or ‘in-context learning’. The correlation between distance in the grid and distance between the representations increases with the length of a random walk on a new graph, but is high from the outset, suggesting the model can rapidly align new stimuli to its learned map (Figure 7h).

## 3 Discussion

We set out to address three questions: how episodic memory shapes semantic memory, how semantic memory shapes episodic memory, and how the two systems interact to provide complementary contributions to memory and problem solving. We hypothesise that semantic knowledge emerges from learning to reconstruct episodes; that episodic memory, which must use neocortical representations for input and output, is compressed using semantic knowledge and therefore shares its biases; and that specific episodic information is combined with general semantic knowledge in working memory for memory retrieval and problem solving. We propose that neocortical memory takes the form of a generative network, trained to reconstruct sequential events replayed by the hippocampus. This enables the network to acquire generalisable semantic knowledge through consolidation, to reconstruct episodes efficiently from highly compressed hippocampal traces, and eventually to capture specific events. During recall, the hippocampus retrieves episodic details which serve as context for the neocortical network to generate predictions based on semantic knowledge. Over time, consolidation allows the neocortex to reconstruct memories independently, reducing reliance on hippocampal traces. Likewise, episodic memories guide problem solving by providing relevant context from the hippocampus for neocortical generation.

In our simulations the neocortical generative network interacts with stored sequences retrieved from the hippocampus as retrieval-augmented generation, but with the neocortex gradually trained on memories from the hippocampus through consolidation, and the hippocampus storing memories in a highly compressed form which can be decoded by the neocortex. Furthermore the stored traces in the hippocampus can combine surprising details with gists in vector form, binding together information at different levels of abstraction in memory.

This model suggests an explanation for several features of human cognition, such as the bidirectional influences between episodic and semantic memory; semantic memory is acquired by learning to reconstruct episodic memories, but also shapes the representation of these episodes, both in terms of their initial encoding, as reflected by the compressed representations observed in hippocampus (*45*), and their gradual schematisation (*20*). To support this, we demonstrated how distortions arise in narratives due to their encoding in compressed form, then increase with consolidation and subsequent forgetting, and compared changes in memory over time to human data (Section 2.1). We also showed how memory reflects priors in the generative model derived from previous experience (Section 2.2). Our model also explains how a number of capabilities improve through consolidation, in addition to the memorisation of ‘replayed’ sequences. In particular, the neocortical generative network supports problem solving, e.g. in relational inference tasks, by learning an implicit structure underlying observations (Section 2.3).

In recent years, neuroscience has seen a move from a modular view of many semi-independent networks learning particular tasks to a focus on the learning of multipurpose representations. Similarly, there has been a transition in machine learning from task-specific models to larger task-general ones trained through self-supervised learning, sometimes referred to as ‘foundation models’ (*24*). The use of a self-supervised task - simply the prediction of the next item in the sequence - to train such models is key to their scaling, as external supervision is not required. We can think of the brain as learning neural ‘foundation models’ too, and this paper suggests how memory consolidation could contribute to their development (see also (*27*)). This view aligns with the growing consensus that prediction error minimisation is key to biological intelligence (e.g. (*80*)), neuroimaging evidence of large task-general networks (e.g. (*81*)), and the relative lack of external supervision in learning. We do not argue that modern LLMs mirror the brain, but that some kind of neural ‘foundation model’ being trained to minimise prediction error on replayed sequences, and used in conjunction with specific memories, provides a good model for a wide range of data on memory, inference and planning. See Figure 8 for a summary of the model’s predictions.

**Figure 8:**
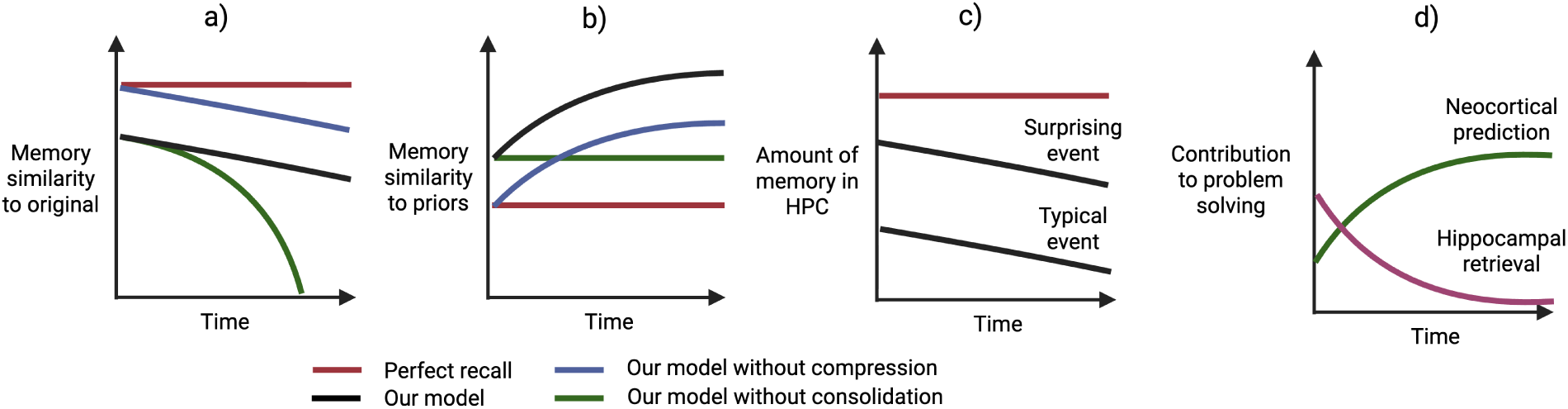
A schematic of our model’s predictions compared to alternative baselines (considering our model with hippocampal forgetting, which is not simulated in the main results). Our model predicts distorted memory from the outset, with the recalled version decreasing in similarity to the original experience (a) and increasing in similarity to priors (b), and compressed encoding with hippocampal contributions to memory decreasing over time (c). Compressed encoding within hippocampus, through use of neocortical representations, explains distortions in newly encoded memories, while consolidation into neocortex ex-plains the increasing similarity to prior experience over time. Furthermore, memories of typical events with well-learned neocortical schemas can be compressed more than surprising events (c). Neocortex and hippocampus collaborate in novel problem solving, with neocortical contributions increasing and hippocampal contributions decreasing with consolidation (d).

### 3.1 Limitations and future work

This work is primarily a model of psychological, rather than neural, data. More could be done to bridge the gap between these ideas and the growing understanding of sequence representations at a neural level. For example, the model allows consideration of navigation, which is associated with distinctive cell types such as place, grid and head-direction cells, so the connection between these different levels of explanation could be explored (see (*22, 82–85*)). There is a trade-off between the complexity of the behaviour that can be simulated and our ability to understand the inner workings of the model. However, progress in ‘mechanistic interpretability’ (e.g. (*86*)) enables the hidden layer activations of LLMs to be compared to neuroimaging data, allowing future comparison of representations at a population level.

LLMs are trained on a simple next item prediction task (*23*), mirroring how prediction error minimisation is thought to drive biological learning (*80*). But even if the objective is shared, the implementations may be very different. In particular, error backpropagation - which calculates each parameter’s ‘responsibility’ for the network’s error - is thought to be implausible because of its non-local nature (*87*). More biologically plausible approximations to error backpropagation do exist (*87*), as well as other families of network that mimic the brain more closely (*80, 88, 89*), which could be explored in future work.

We draw on a body of research connecting memory and schema formation (*8, 12, 90*). The SLIMM framework (*8*) suggests that whilst high prediction error memory elements may be preferentially encoded in hippocampus, schema congruent information may be remembered more easily by cortex. Consistent with this account, our model could predict a U-shaped relationship between surprise and recall performance, in which moderately surprising elements - those not unpredictable enough to be encoded in detail, nor familiar enough to be captured by existing schema - are especially prone to forgetting (see (*91*)). Whilst we show that memories become more schema congruent overall in our simulations, and propose a mechanism for the storage of surprising details in hippocampus, we do not model how the retention of different aspects of a memory depends on their predictability, and how this changes over time. Future research could compare our model’s predicted pattern of memorability as a function of schema congruence with these data.

Memories bind together neocortical representations, with anterior hippocampus connected to anterior neocortical regions capturing more conceptual content, and posterior hippocampus connected to posterior neocortical regions capturing more sensory content; consolidation is associated with growing dependence on the former (*46*). If the gist is associated with anterior neocortex, future work could test the prediction that the more schema congruent an episodic memory, the greater the dependence on anterior as opposed to posterior hippocampus.

A related issue that could be explored further is how the hippocampal encoding evolves over time. In our model, hippocampal traces are assumed to fade once the neocortical network can approximately reconstruct an event, but this is not simulated explicitly. A complexity of this process is that the stored gist representation must be ‘in sync’ with the neocortical model to be decompressed correctly, so as neocortex changes the gist would drift from its original meaning, which could be another factor leading to hippocampal forgetting. Long-term retention of hippocampal traces might therefore require some mechanism to preserve the correspondence between conceptual representations in hippocampus and in cortex (*92*). A separate question is whether different elements of a hippocampal trace decay uniformly, or whether details might fade away once they become more predictable, with the gist representation the last element to be forgotten.

Whilst experience is continuous, it is discretised into events in memory (*93, 94*), a process known as event segmentation. Event segmentation is thought to be based on prediction error (*95–97*) (but see (*98*) for an alternative account). Whilst the sequences used in our simulations are ‘pre-segmented’ for simplicity, this is potentially consistent with our model, as the generative network can provide an ongoing measure of ‘surprise’ during perception (*40, 96, 97, 99*), e.g. as quantified by perplexity (*23*). Future work could explore the connections between these ideas.

Human learning can display leaps of insight (e.g. (*100*)), with abrupt improvements thought to correspond to the reorganisation of mental representations (*101*). Future work could explore the development of an LLM’s internal representations as a task is learned, comparing this to the corresponding ‘behavioural’ changes, to see if sudden increases in performance correspond to sudden representational changes. This could then be compared to the dynamics of learning in humans.

Future research could also explore how retrieval, encoding, and replay processes are orchestrated. We simplify matters by using pre-segmented sequences, encoding every event with a fixed level of detail, and ‘hard-coding’ when the hippocampus is queried. However, one could train a ‘meta-controller’ to co-ordinate these processes with reinforcement learning (*102*). This might learn to use different cognitive resources when it was optimal to do so, e.g. prioritising memories for replay to maximise future performance, and only encoding memories when it is worth the cognitive cost. This could build on previous studies like (*103*), which prototypes these ideas with simpler models.

Finally, catastrophic forgetting refers to the overwriting of old knowledge by new knowledge when a neural network learns multiple tasks or distributions consecutively (*104*). Whilst gradual forgetting is expected in cognition, catastrophic forgetting is a more dramatic interference of new with old knowledge, which is not typically observed. The LLM’s gradient of forgetting in Section 2.1.2 is steeper than would be expected of human memory, suggesting that the brain has mechanisms for maintaining neocortical memories over the course of a lifetime that LLMs do not. Future research could investigate these mechanisms, e.g. whether generative replay in neocortex might facilitate continual learning (*92, 105*).

### 3.2 Episodic and semantic memory

In our model episodic and semantic memory combine to reconstruct the past and predict the future, but how clear is this distinction? If a series of narratives representing ‘episodes’ are ‘consolidated’ into the neocortical LLM, the resulting LLM could support both memory for specific episodes (*28*) and for semantic ‘facts’, with the latter learned as a side-effect of reconstructing the former. Not only are ‘beliefs’ influenced by ‘episodes’ the network was trained on, but the ‘episodes’ are reshaped by the ‘beliefs’, as we saw in a range of memory distortion simulations. This can be the case even for the initial hippocampal encoding of episodes in the full model, as demonstrated in Section 2.1.

How do we distinguish remote memory from imagination, if both consist largely of schema-based predictions from a generative model? The generative model’s prediction error for sequences from the training data tends to be lower than for novel sequences (*106*), so the generative network could provide a measure of recognition memory even if the original hippocampal trace has faded away. However this might correspond to a ‘sense of familiarity’ rather than definitive knowledge that a stimulus was experienced. Thus, the presence of similar real memories (*61, 62*), or the rehearsal of imagined events (*107*), can induce false memory, presumably from a ‘sense of familiarity’ in the generative network. This could align with hippocampal amnesics’ sometimes preserved ability to make relative judgements of familiarity (*108*).

On the other hand, episodic memories arguably need to include some details from a hippocampal trace to be recognised confidently (*108, 109*). Memories may retain some limited trace in the hippocampus proper for a long time, so the retrieval of event-unique details which have not yet been consolidated could help to distinguish real memories from imagination. The neocortical network may contain many semantic facts relating to one’s personal history, e.g. that one went on a holiday to a particular place as a child, and could thus construct events consistent with these facts. However, if episodic recollection requires retrieval of unique details encoded during the event, then even an accurate reconstruction might be considered a ‘semanticized’ memory rather than a truly episodic one (e.g. (*7*)).

### 3.3 Conclusion

We have proposed a model of the bidirectional interactions between episodic and semantic memory, in which semantic memory in neocortex relies on a generative network, trained to reconstruct sequential events replayed by the hippocampus. Episodic memories are encoded in the hippocampus in compressed form optimised for efficient neocortical reconstruction, and recalled by ‘decompressing’ these traces with the generative network. In other words, hippocampus and neocortex work together in encoding, recall, and problem solving to provide episodic details and predictions from semantic knowledge based on these details respectively. During consolidation, these reconstructions are then replayed to update the neocortical generative network to predict the next item in each sequence. We make use of recent advances in the large language model literature to simulate this hippocampo-neocortical interaction as ‘retrieval-augmented generation’ (RAG), with the addition of mechanisms to *compress* episodic memories and to *consolidate* them into the generative network. This model explains changes in the characteristics of autobiographical memories over time, and accounts for gist or schema-based distortions. As well as capturing the gist of specific episodes, the neocortical network extracts statistical patterns that generalise to new situations. We also show how the hippocampal and neocortical networks could jointly contribute to problem solving through RAG, allowing memories retrieved from the hippocampus to provide the context for prediction using the ‘general knowledge’ of the neocortical network. We demonstrate that, by simply learning to predict the next item, the neocortical network can acquire abstracted knowledge of commonalities across experience, which can be used to support inference and planning.

## 4 Methods

### 4.1 Large language models

Our simulations use large language models (LLMs), which are neural networks trained to predict the next item in a sequence based on preceding items (*23*). Most commonly, the sequence is text and the items are small chunks of characters (or ‘tokens’). These models operate autoregressively: given a partial sequence, they estimate a probability distribution over possible next items. Training consists of adjusting the network’s parameters so that it assigns high probability to the actual next item observed in the data, a procedure known as causal language modelling.

Once trained, the model can be used to generate new sequences by repeatedly predicting and selecting the next item. The simplest approach is greedy decoding, in which the most probable next item is always selected, but sequences can also be generated by sampling from the model’s predicted probability distribution, with a temperature parameter controlling how variable the output is. Higher temperatures increase variability by flattening the distribution, while lower temperatures favour the most likely continuations.

Most LLMs use a mechanism called attention (*110*), which allows each element to be enriched by ‘attending to’ information earlier in the sequence. This matches the intuition that the meaning of a word is understood in the context of the text so far, not in isolation as in earlier approaches to text processing with machine learning.

See Section S1.1, SI, for further details.

### 4.2 Modelling hippocampal-neocortical interaction as retrieval-augmented generation

#### 4.2.1 The compression and encoding of episodic memory

In Sections 2.1 and 2.2 we use Mistral-7B-Instruct-v0.2 (*29*) to explore the compression, encoding, retrieval-augmented generation, consolidation, and forgetting of narrative events. In addition to the LLM itself, pre-trained weights from (*33*) are used to transform episodes into compressed xRAG vectors. In brief, these weights are trained so that the embedding of the text reconstructed by the LLM is as close as possible to the embedding of the original. A key advantage is that the underlying idea in (*33*) is not specific to language but could in principle work for arbitrary types of sequence. Note that xRAG weights are only compatible with a particular LLM; Mistral-7B-Instruct-v0.2 is used because the xRAG weights are trained to transform embeddings into token representations of this specific model. But in principle the context compression mechanism could work for any LLM. See Section S1.1.4, SI, for a discussion of alternative context compression approaches.

To identify surprising details given the xRAG vector, we first segmented the original story into short clause-like phrases using the spaCy Python library, splitting at (i) sentence boundaries and (ii) commas or conjunctions. To quantify how *unpredictable* each candidate phrase 𝑝 is given the gist vector, we scored it by conditional perplexity. Specifically, we concatenated the gist vector with the candidate phrase and computed the model’s perplexity. Higher perplexity indicates that the phrase is less predictable from the gist. For a requested detail level 𝑛, we selected the top-𝑛 highest-perplexity phrases as the ‘unpredictable details’.

The prompt used to recall a memory with retrieval-augmented generation is as follows, formatted as suggested in the xRAG (*33*) code. Note that the ‘INST’ tag denotes the instruction for Mistral 7B, which is an instruction-tuned model:

[INST] Refer to the background document and answer the question.

Background: *{*XRAG TOKEN*}*

Question: *{*FIRST LINE*}* What happened (in detail)?

Other details to include: *{*SURPRISING DETAILS*}*. [/INST] The answer is:

Here ‘XRAG TOKEN’ is the single token encoding the gist produced by the xRAG projection weights, and ‘SURPRISING DETAILS’ is a comma-separated list of the *n* highest perplexity phrases. (The line beginning ‘Other details’ is omitted from the prompt when no details are provided.)

#### 4.2.2 Modelling the hippocampus

Recall requires both pattern completion of the current item and prediction of the next one. For example, recalling a day out from a photo requires first retrieving the scene depicted in the photo, and then remembering the events following and preceding that scene. Modelling sequence memory therefore requires a mechanism that supports both autoassociative reconstruction of the present state from a partial input and heteroassociative transitions to future states (*111*).

Here this is implemented with a pair of modern Hopfield networks (see Section S1.1.5, SI, for further details). A standard modern Hopfield network (MHN) retrieves items from their noisy versions (*31*), while an asymmetric MHN (*112*) retrieves the next item for a given item. This simple model demonstrates the idea of combining autoassociative and heteroassociative connectivity in a ‘biologically plausible’ way (*113*), but many alternative implementations of sequence memory exist, e.g. (*114*), and could be compatible with this proposal. In our model, the xRAG gist vector for an event can be retrieved autoassociatively from a cue, while each token in the sequence of details can be retrieved heteroassociatively from the previous token. (Note that we consider transitions going forward in time for simplicity, but a fuller model could involve heteroassociative transitions going backward in time too.)

To encode text as sequences of characters, it is crucial to distinguish between different instances of the same character, i.e. the correct next character after ‘a’ depends on the letters that precede ‘a’. (Because the same words feature in different episodes, the history that determines the correct transition stretches back many characters.) The asymmetric MHN is by default a ‘Markov chain’ model of sequential memory, so it is necessary to use a state representation that captures the history. This is based on previous work involving the role of temporal context in memory (*36, 115*). Here we simply add a decaying representation of the previous item to a given item’s representation, with the weighting of the previous item set by a decay constant.

In summary, each element of a sequence is represented by a state vector, which consists of the current state plus a decaying trace of previous states. In the main text simulations, the first such state is the xRAG vector, and subsequent states are characters in the string of unpredictable details. (In Section 2.3, which omits the compression aspect for simplicity, the states are characters in the string.) To describe this in terms of the Krotov and Hopfield (*113*) formulation, the values of each state vector are encoded in the weights to and from a given memory unit in an MHN, while the asymmetric MHN stores state 𝑛 in incoming weights to a memory unit, and state 𝑛 + 1 in outgoing weights. Retrieval proceeds in two stages (described with reference to the main text simulations). First, given a query xRAG vector computed from the query text, the MHN computes similarities to stored states (Figure S1a), and applies a softmax function with inverse temperature 𝛽. In the limit of high 𝛽, this results in autoassociative pattern completion of a single stored state, ideally that of the stored gist vector. Subsequent elements, corresponding to characters of the unpredictable details, are then retrieved heteroassociatively, with the output of one retrieval becoming the input to the next. A range of values of the decay constant described above can give good results (Figure S1b). High 𝛽 is also used here to retrieve a single next state prediction rather than a superposition (Figure S1c).

Note that we do not model forgetting in the hippocampus, or errors that result from retrieval failures rather than compressed encoding. The ability to retrieve stored sequences is near perfect for the parameters chosen, as shown in Figure S1, but future work could explore how retrieval errors in the hippocampus affect the behaviour of the system.

#### 4.2.3 Consolidation of reconstructed episodes

In simulations involving Mistral 7B, the model is fine-tuned in a different way from the ‘usual’ end-to-end method because of its size. Instead of updating the entire model, parameter-efficient fine-tuning (PEFT) methods update only a small subset of parameters. One such method is low-rank adaptation (LoRA; (*68*)), which injects trainable low-rank matrices alongside frozen pre-trained weights. We implement LoRA with the ‘PEFT’ Python library (*116*).

We used a LoRA rank of 𝑟 = 16, scaling parameter 𝛼 = 16, and dropout probability 0.05. LoRA adapters were applied to the linear weight matrices used to compute the query, key, value, and output projections in the attention mechanism, as well as to the gate, up, and down projection matrices of the feed-forward network.

The training data is formatted as follows, in the form recommended for an instruction-tuned model such as Mistral 7B:

[INST] *{*FIRST LINE*}* What happened (in detail)? [/INST] *{*REST OF STORY*}*

The prompt used to test *consolidated* episodic memory, i.e. memory without any retrieval from the model hippocampus, is as follows:

[INST] *{*FIRST LINE*}* What happened (in detail)? [/INST]

The prompt used to test *consolidated* semantic memory is as follows, where the prompt contains enough context to pick out a single relevant story, and the question has an answer in that story:

[INST] Remember the story in which *{*FIRST LINE*}*. *{*QUESTION*}* Answer with a word or short phrase. [/INST]

#### 4.2.4 Analysing the HIPPOCORPUS and NFRD datasets

The HIPPOCORPUS dataset (*18*) consists of 2779 recalled stories, 1319 retold ones, and 2756 imagined ones (not included in these analyses). The Naturalistic Free Recall Dataset (NFRD; (*19*)) consists of 116 recalls for the ‘pieman’ story, 116 for ‘eyespy’, 113 for ‘pieman’, and 113 for ‘oregontrail’.

The following prompt was used to label the attributes of recalled and retold items in the HIPPOCORPUS dataset, and of original and recalled items in the NFRD dataset with gpt-4o-mini via the OpenAI API:

Your task is to score text on three metrics: how concrete (vs abstract) it is, how rich in detail it is, and how specific (vs general) it is.

Return ONLY a JSON dictionary with 3 keys, each a float 0-1:

*{*‘concrete vs abstract’: 0–1, ‘rich vs poor details’: 0–1, ‘specific vs general’: 0–1*}*

A higher score corresponds to more concrete, richer in detail, or more specific text.

This reflects standard practices for using LLMs for data annotation (*117*).

### 4.3 Modelling the interplay of semantic and episodic memory in consolidation

Event extension and contraction are modelled with simple stories in text form, each only a few lines long (*72*). Three types of narrative are used. Some are ‘typical’ stories, unmodified from the (*72*) dataset, whereas an ‘incomplete’ story has characters removed, and an ‘updated’ story has characters added, so it continues beyond its natural ending. Note that this is a challenging task as some stories could be interpreted as either incomplete or updated. To disentangle the effect of story category from the effect of length, the incomplete and updated stories were all the same length. To achieve this, text from a different random story was added to stories shorter than average, extending them to the mean length, while longer than average stories were shortened to the mean length.

After encoding and reconstructing the stories, the Mistral 7B model (*29*) was fine-tuned on the reconstructions for three epochs and then recall was tested by giving the model the first 150 characters and observing the output.

### 4.4 Neocortical learning of sequential structure for inference and planning

In Section 2.3 we train sequence models (GPT-2; (*23*)) on non-linguistic sequences, choosing a smaller model to run more extensive simulations of learning structural inference from scratch. The ‘medium’ sized GPT-2 model is used, which has 345 million parameters including 24 transformer blocks, and is trained from scratch with randomly initialised weights. See Table 5 for a summary of the models trained.

**Table 5:**
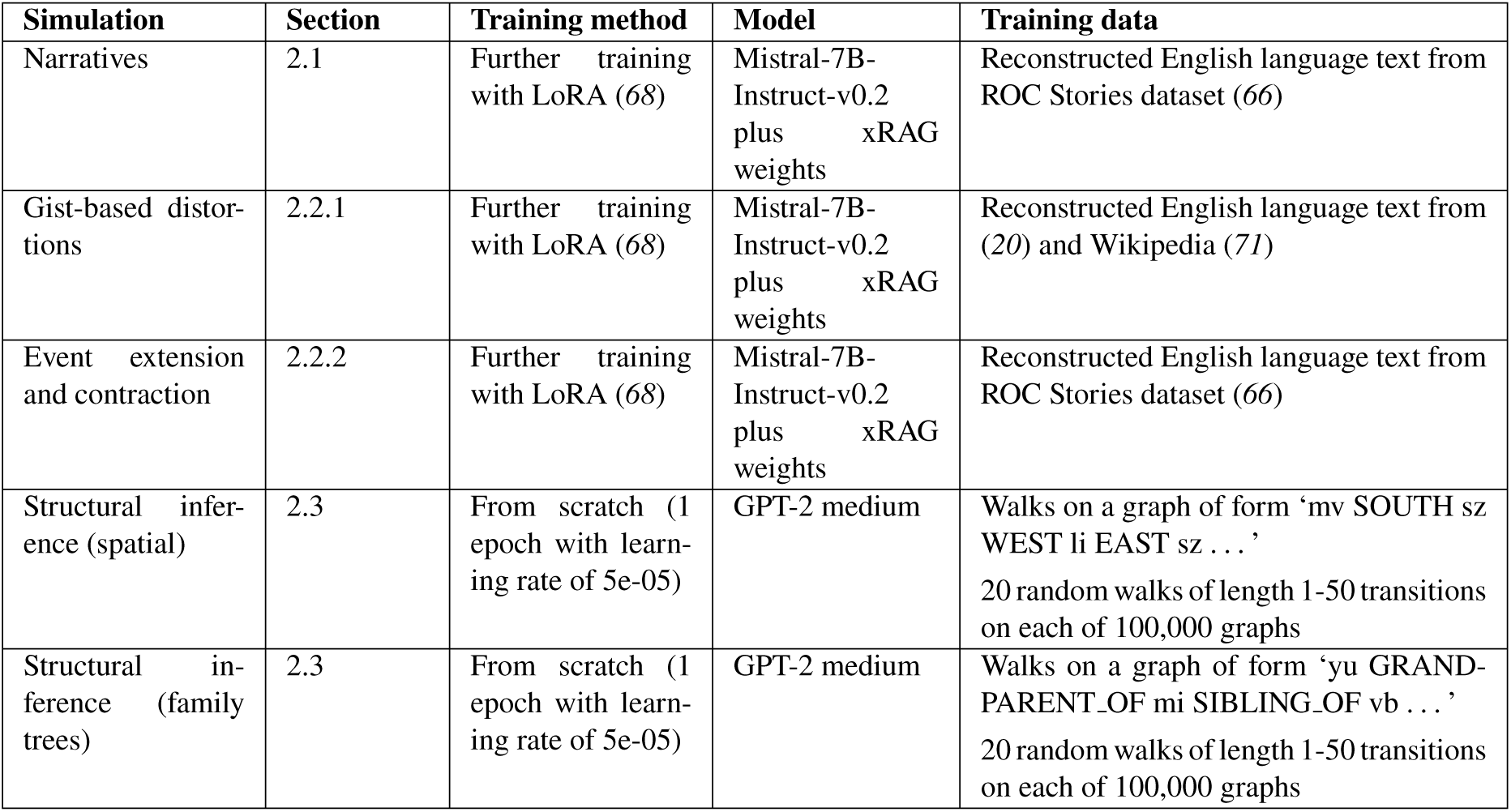
Summary of models trained. Here ‘reconstructed’ means the version of the original data reconstructed by the LLM from its compressed encoding.

| Simulation | Section | Training method | Model | Training data |
| --- | --- | --- | --- | --- |
| Narratives | 2.1 | Further training with LoRA (68) | Mistral-7B-Instruct-v0.2 plus xRAG weights | Reconstructed English language text from ROC Stories dataset (66) |
| Gist-based distortions | 2.2.1 | Further training with LoRA (68) | Mistral-7B-Instruct-v0.2 plus xRAG weights | Reconstructed English language text from (20) and Wikipedia (71) |
| Event extension and contraction | 2.2.2 | Further training with LoRA (68) | Mistral-7B-Instruct-v0.2 plus xRAG weights | Reconstructed English language text from ROC Stories dataset (66) |
| Structural inference (spatial) | 2.3 | From scratch (1 epoch with learning rate of 5e-05) | GPT-2 medium | Walks on a graph of form ‘mv SOUTH sz WEST li EAST sz . . . ’<br>20 random walks of length 1-50 transitions on each of 100,000 graphs |
| Structural inference (family trees) | 2.3 | From scratch (1 epoch with learning rate of 5e-05) | GPT-2 medium | Walks on a graph of form ‘yu GRAND-PARENT_OF mi SIBLING_OF vb . . . ’<br>20 random walks of length 1-50 transitions on each of 100,000 graphs |

We simulated how consolidation enables structural inference by the neocortical network, considering the example of a spatial graph and a simple family tree graph (as in (*22*)). We trained GPT-2 medium from scratch on random walks on these graphs, corresponding to sequences of observations encoded in the hippocampus, and then tested the models’ inference abilities on novel sequences with the same underlying structure.

Note that on some of the family tree inference problems there are multiple possible answers which are consistent with, but cannot be inferred from, the sequence so far (such as the imagined family member ‘ef’ in ‘ab CHILD OF cd PARENT OF ef’, whereas ‘ab’ is the expected answer), and these are counted as incorrect, making this quite a harsh performance metric.

We simulate episodic contributions to problem solving as retrieval-augmented generation, using the two models in the structural inference results above (one trained on spatial graphs, and one on family tree graphs, such that each has learned the structural regularities of the stimuli). In each case, the hippocampus encodes sequences from *new* spatial or family tree graphs, corresponding to observations in a new spatial environment or of a new family’s relationships. 200 new graphs were constructed for each task, each missing a randomly chosen edge. Two short walks on each graph, which did not include the missing edge, were stored in the ‘hippocampus’ (simply a list of strings in this example). The two walks were chosen so that solving the task requires combining information from *both*, e.g. it requires both ‘ab EAST cd SOUTH ef’ and ‘ef WEST gh’ to infer that ‘gh NORTH ab’. For each missing edge, a query of the form ‘gh NORTH’ or ‘cd PARENT OF’ was constructed for the spatial and family tree graphs respectively. Testing involved two stages, retrieval followed by generation: first the hippocampus is queried for relevant traces, simply by finding sequences from the same environment or family tree. Then the generative network produces an output conditioned on the retrieved sequences concatenated with the sequence for the task (Figure 6j), i.e. with the retrieved sequence and the current task in the LLM’s prompt. Greedy decoding was used to generate predictions.

## Acknowledgments

We thank Oliver Vikbladh, Misun Kim, Steve Fleming, Kris Jensen, and Mathias Sablé-Meyer for their insightful comments on the manuscript.

## Author contributions

Conceptualization: ES, NB. Methodology: ES, NB, Investigation: ES. Visualization: ES. Supervision: NB. Writing: ES, NB.

## Data and materials availability

Code for all simulations can be found at https://github.com/ellie-as/hippocampal-neocortical-RAG. All datasets used are publicly available.

## S1 Supplementary information

### S1.1 Further model details

#### S1.1.1 Modelling sequence learning

The simulations described above use autoregressive sequence models to represent the generative networks trained through hippocampal replay. Specifically transformer-based deep neural networks for text generation are used.

The primary goal during training is to adjust the model’s parameters through maximum likelihood estimation, so that the probability it predicts for the true next item in each sequence, based on the items so far, is as high as possible. In other words, the network’s weights are updated to predict the probability distribution of the next item as accurately as possible, a process known as causal language modelling. The training data for the original GPT-2 model is the WebText dataset of online content. Once the model is trained, it predicts the probability distribution across all items given the items so far. To generate a sequence one can sample from this distribution or simply take the most probable item at each step. The equation below gives the probability of a sequence 𝑥 as a product of conditional probabilities of its items:

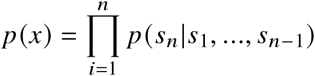

We use the Transformers Python library (*118*) in all simulations. The first stage of using an LLM is to prepare the inputs with tokenisation. A tokeniser maps commonly occurring chunks of characters to IDs (in order to look up the right token embedding in a learned embedding matrix); in the case of language tokens are often words or parts of words. The concept of tokenisation is applicable to arbitrary sequences, but for simplicity and consistency across the simulations all stimuli are converted to strings of characters (if they are not already text-based). For Mistral 7B the same tokeniser is used as in the pre-trained model, but for the inference models a custom one is used, to ensure that relationships (e.g. EAST or CHILD OF) and entities (e.g. ‘xy’) were each represented by a single token. (Without this, the default tokeniser does this inconsistently, e.g. with some pairs such as ‘be’ represented as one token because commonly co-occur, and other such as ‘bx’ as two tokens because they do not.)

What exactly does causal language modelling with a custom dataset involve? First the training data is split into blocks, and then for every block the cross-entropy loss is aggregated across all the next token prediction tasks within the block. For example, for GPT-2 the block size (which is also the context size of the trained model) is 1024 tokens. This means that the model is trained to consider up to 1024 tokens of context when predicting the next token in a sequence. So for each block the model tries to predict the second token based on the first token, then the third token based on the first two, and so on, until it predicts the final token based on the preceding 1023.

The loss measures the difference between two probability distributions: the distribution predicted by the model and the actual distribution in the data. For each token prediction task, the actual distribution is a ‘one-hot’ vector with a one for the real next token and zeros elsewhere. Specifically, the cross-entropy loss for a single prediction task is calculated as the negative log probability assigned by the model to the actual next token. For a block of tokens, the total loss is the sum of the cross-entropy losses for each token prediction task within the block, and the weights of the model are updated based on this total loss. This procedure is the same whether the model is fine-tuned or trained from scratch.

The learning rate decreases over the course of training in all simulations, as per the default behaviour of the Trainer class in the Transformers Python library (*118*).

#### S1.1.2 Sampling options

There are many ways to generate sequences from a trained sequence model like Mistral 7B or GPT-2. The simplest is greedy decoding, where the model selects the token with the highest probability as the next token in the sequence. However this can lead to repetitive and predictable sequences. Sampling from the learned probability distribution with a given temperature introduces randomness into the selection of the next item, and provides a way to control the model’s ‘imaginativeness’. Specifically, the temperature parameter determines the ‘sharpness’ of the distribution from which output tokens are selected, so that sequences at a higher temperature are more ‘imaginative’, but more likely to be nonsensical.

The equation below describes the ‘softmax with temperature’ function that is applied to the vector of scores for each token to produce a vector of *probabilities*. A large temperature T flattens the distribution, whereas T close to zero approximates a ‘one-hot’ vector, with a probability of one for the most likely token:

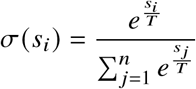

Unless stated otherwise, sequences are sampled from the probability distribution of tokens with a given temperature. Beam search is a search strategy used in some simulations, which considers multiple potential paths through the model’s probability ‘landscape’. At each step, it keeps a fixed number (the beam width) of the most probable sequences generated so far and extends them, eventually choosing the sequence with the highest overall probability.

#### S1.1.3#Further details on attention

In short, the attention mechanism works as follows (explained with reference to GPT-2): a query, a key, and a value vector are produced for each element in the sequence by learned weights. For a given element, the model computes a set of attention scores which determine how much attention should be paid to each element of the input sequence when representing this element. This is achieved by taking the dot products of the given element’s query vector with the key vectors for each element. The final output is the sum of the value vectors for each element weighted by the corresponding attention scores (with some normalisation), allowing the model to aggregate information from the most relevant parts of the input. This describes a single ‘attention head’, but GPT-2’s architecture features multiple attention blocks, each with multiple attention heads, which tend to develop different ‘specialisms’ through training. To be more precise the equation is given below:

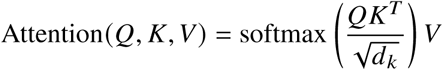

In this equation, 𝑄, 𝐾 and 𝑉 represent the query, key, and value matrices, respectively. These are derived from the input data. 𝑑_𝑘_ represents the dimensionality of the keys (and queries), and is used to scale the dot products in a way that leads to more stable gradients. The softmax function is applied to ensure the weights sum up to 1. When the resulting term is matrix multiplied by V, the result is the relevance-weighted sum of the vector representations in V, for each token in V. In other words, the attention mechanism enriches a token’s representations with a relevance-weighted sum of the other tokens’ representations. See (*119*) for a detailed description of this process.

Self-attention is a special case of attention in which elements of a sequence ‘attend to’ the sequence they are a part of. Specifically, ‘masked’ self-attention is used, meaning that the representation of a token in an attention block only ‘attends to’ preceding tokens. Previous models like BERT (*120*) perform much less well at text generation partly because their attention is not masked in this way.

Transformer blocks come in different varieties. GPT-2 (*23*) is a decoder-only model consisting of a stack of transformer decoder blocks. Each such block consists of a masked self-attention layer followed by a feedforward layer. In a nutshell, inference with GPT-2 works as follows: First an embedding for each token is obtained from a learned embedding matrix. A key point is that position embeddings representing the tokens’ positions in the sequence are added to the token embeddings (these are required because self-attention is permutation-invariant, so otherwise information about the order of items in the sequence would be lost). Then the sequence of embeddings passes through a series of transformer decoder blocks, each further enriching the token representations so that they come to capture the meaning of the text. See (*121*) for helpful illustrations of the stages of processing in GPT-2. GPT-2 (*23*) comes in several sizes, with the number of transformer blocks depending on the size of the model. The small variant has 12 transformer blocks, the medium variant has 24 transformer blocks, and the large variant has 36 transformer blocks.

#### S1.1.4#Context compression comparison

Context compression has become an important problem in LLM research because performance often depends on access to relevant information, but context size is limited. Even when long-context models are available, this is computationally costly, and providing too much distracting information to the model can degrade performance.

Context compression techniques for LLMs can be divided into extractive and abstractive approaches, where the former selects a subset of the original context, and the latter summarises the context in some way. Extractive methods include LLMLingua (*122*), in which an auxiliary model predicts the importance of each token for a given LLM, and only the most important tokens are included in the context.

One intuitive form of abstractive context compression simply produces a textual summary of the input, e.g. see RECOMP (*123*). But abstractive methods can also map long contexts into compact latent representations. The in-context autoencoder approach (*63*) compresses arbitrarily long inputs into a fixed number of memory slots. The encoder’s weights are trained in such a way that the LLM can reconstruct the original context conditioned only on the memory slots. Intriguingly, the authors find that the bigger and better the LLM, the more the context can be compressed at the same reconstruction accuracy.

The xRAG (*33*) method used here is similar to ICAE, but compresses the context to a single token. The primary reason for using xRAG rather than ICAE is because the stimuli in our simulations are short, and the pre-trained ICAE encoders are designed to compress contexts to 128 tokens, which can be longer than the stories themselves. However there is no strong theoretical reason to prefer xRAG to other candidate methods for abstractive context compression. One drawback of xRAG is that compressing texts to a single token then using many tokens to encode details is odd, as the core of the memory takes up a fraction of the capacity of the more extraneous details.

#### S1.1.5#Modelling sequence memory in the hippocampus

Here we provide further context to motivate the model of the hippocampus.

In the static model in (*27*), a modern continuous Hopfield network (MCHN; (*31*)) represents the hippocampus, interpreted such that the feature units activated by an event are bound together by a memory unit (*113*). In the sequential model, this could be adapted to store sequences as follows. Let us consider sequences represented as strings of characters or symbols, which can capture language, spatial trajectories, transitions in a graph, stimuli in sequential learning tasks, and more.

(*58*) give a unifying account of how neural network models of associative memory such as MCHNs operate, which helps to explain the extension to sequences. They develop the biologically plausible modern Hopfield network (*124*) into ‘a general framework for understanding the operation of…memory networks as a sequence of three operations: similarity, separation, and projection’, which they term ‘universal Hopfield networks’ ((*58*), Abstract). They observe that any of these models can be made asymmetric or heteroassociative rather than autoassociative, if the projections from feature units to memory units capture the current state, but the projections from memory units back to feature units are different (for sequences, these projection weights would correspond to the ‘next state’). Then one state retrieves the next, rather than each state retrieving a pattern-completed version of itself. That is, we want to store patterns 𝑋 and ‘next patterns’ 𝑋_𝑛𝑒𝑥𝑡_ instead of just patterns, and change the MCHN’s update rule accordingly:

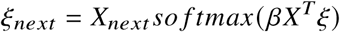

The modern continuous asymmetric Hopfield network (MCAHN) converts the MCHN to work for sequences, consistent with the framework above (*112*). However there is a problem when you try to apply this naively to certain sequences, as the MCAHN is by default a ‘Markov chain’ model of sequential memory. Whilst it works for sequences in which items are unique, it doesn’t work for, e.g., sentences without modification, as it cannot accurately remember sequences with repeated items; see Figure 5a of (*125*). When an item is repeated in a sequence, (*125*) show that a MCAHN has trouble recalling the next item, instead producing a composite.

(*125*) introduce the temporal predictive coding network (tPC), a more complex network learnt by gradient descent, based on the intuition that higher layers minimize prediction error at lower layers (*126*). (A drawback of the tPC is that the model must be ‘presented with the sequence for multiple epochs’ until convergence, so it is no longer capable of one-shot learning, which arguably undermines its plausibility as a model of hippocampal encoding.) They show that the one-layer tPC also has the MCAHN’s issue with repeated items, but suggest the two-layer tPC to resolve this. As (*125*) show in Figure 5d of their paper, the hidden layer represents an item’s position in the sequence so the next item can be recalled correctly.

Alternatively, one could combine the concept of the modern asymmetric Hopfield network (with return projections from memory to feature units representing the next state) with a state representation that captures the history. This is based on previous work involving the role of temporal context in memory (*36, 115*). Consider the example of encoding sentences, i.e. sequences of characters. Suppose each state is a vector of length equal to the number of symbols, consisting of one at the index of the current symbol, plus the previous state multiplied by some decay factor. See Figure 9 for an example. This has the benefit of still being compatible with one-shot learning, although the memory capacity may not scale as well as the more complex predictive coding model approach. Initial testing suggests that this model can encode and retrieve sentences (although retrieval performance goes down with sentence length).

**Figure 9:**
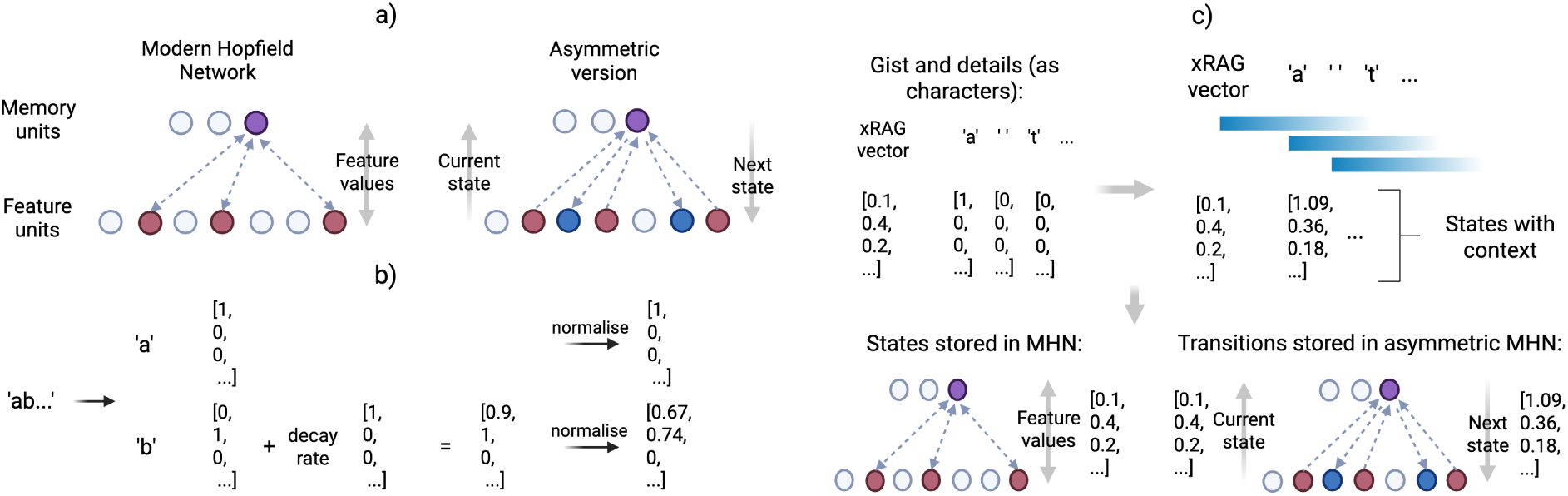
Using modern Hopfield networks (MHNs) to model the hippocampus. a) In asymmetric versions of MHNs, the weights between feature and memory units differ, as visualised from the perspective of (*113*). b) A sequence of arbitrary symbols, e.g. ‘ab…’, is represented as a sequence of vectors over the feature units, where each vector is the sum of the current state plus the decay rate (here set to 0.9) times the previous state. c) To store a narrative in compressed form, the xRAG vector and details are represented as a series of states, with a decaying trace of previous states. The states are encoded in an MHN, and the state-next state transitions in an asymmetric MHN. In the latter, for each state the return connections from memory units to feature units encode the next state.

To show how the preceding ‘context’ is captured, consider the example of encoding sentences, i.e. sequences of characters. Each state can be represented by a vector of length equal to the number of symbols, with one at the index of the current symbol, plus the previous state multiplied by some decay factor. In other words, we want a vector representation **x_i_** of the i^th^ state which is a sum of the current symbol’s vector **v** and activity from the previous state **x**_𝑖−1_ multiplied by a decay rate 𝜆 (noting that the states are then normalised to unit length):

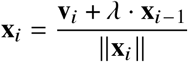

As with all modern Hopfield network (*31*) variants, the inverse temperature, 𝛽, is a key parameter determining the network’s behaviour. In this case a high value of 𝛽 would be desirable to avoid composite states. The softmax function is given below; in the limit of high 𝛽, the output of this function becomes a vector with one at the index of the maximum and value and zero elsewhere:

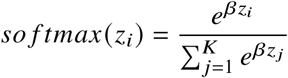

Episodic memories consist of complex representations at each time step, with autoassociative connections thought to reconstruct the current time step, and heteroassociative connections to activate the next (see (*111*)). As described in Section 4.2.2, the asymmetric MHN described above is combined with a ‘standard’ autoassociative MHN for retrieving an item from a noisy query.

Whilst this simple model helps justify the assumption that the hippocampus can store sequences associatively, it clearly has some limitations. For example, it suggests that when a state from a certain point in an encoded sequence triggers recall, states preceding that point cannot be recalled (only the subsequent states). This could be addressed by adding heteroassociative connectivity backwards in time. A more fundamental problem is that the cue for retrieval can be a noisy version of the whole sequence, but in this simple model retrieval begins by denoising a single item in isolation. This means that retrieval may perform poorly when all items are so noisy that retrieval of any item individually is poor, but the sequence as a whole is sufficient to identify the right memory. Future work could investigate biologically plausible models of hippocampal memory that address this.

### S1.2 Supplementary results

#### S1.2.1 Supplementary figures

Figure S1 shows some additional results relating to the model hippocampus.

Figure S2 shows the results of performing principal component analysis on token representations in the spatial inference model, across all layers of the model.

#### S1.2.2 Supplementary tables

Table S1 shows examples of encoded and recalled stories from Section 2.1.

Tables S2, S3, and S4 show that the patterns in concreteness, specificity, and richness in detail hold even when controlling for length. This is important because for both the HIPPOCORPUS and simulated data, there are significant Pearson correlations between length and each attribute (Table S2). For the HIPPOCORPUS data, regressing out the length, and then comparing the residual score for each attribute (i.e. the component which cannot be explained by the length) using a t-test, gives the results in Table S3. Doing the same for the simulated data gives the results in Table S4.

Table S5 shows the Bartlett story recalled at different temperatures, illustrating the approximate memorisation of the narrative in Section 2.2.1. Tables S6 and S7 give the inference accuracy for each template tested in Section 2.3.

---

**Figure S1:**
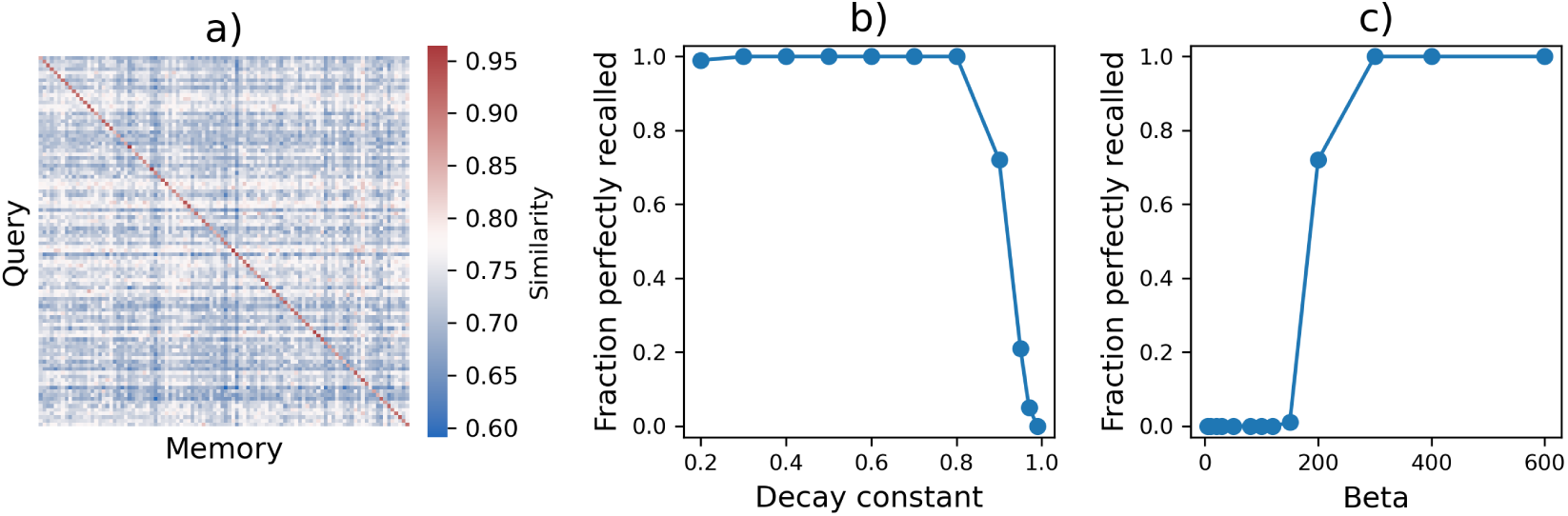
Modelling the hippocampus with a dual-MHN model for 100 stories from (*66*). a) Similarity (dot product) between each xRAG query embedding (constructed from an 80-character excerpt) and the stored xRAG story embedding (whole story), showing a strong diagonal indicating correct memory selection. b) Fraction of episodes perfectly recalled as a function of the decay constant. c) Fraction of episodes perfectly recalled as a function of inverse temperature 𝛽 (applied identically to both MHNs), showing improved recall with sharper retrieval.

**Figure S2:**
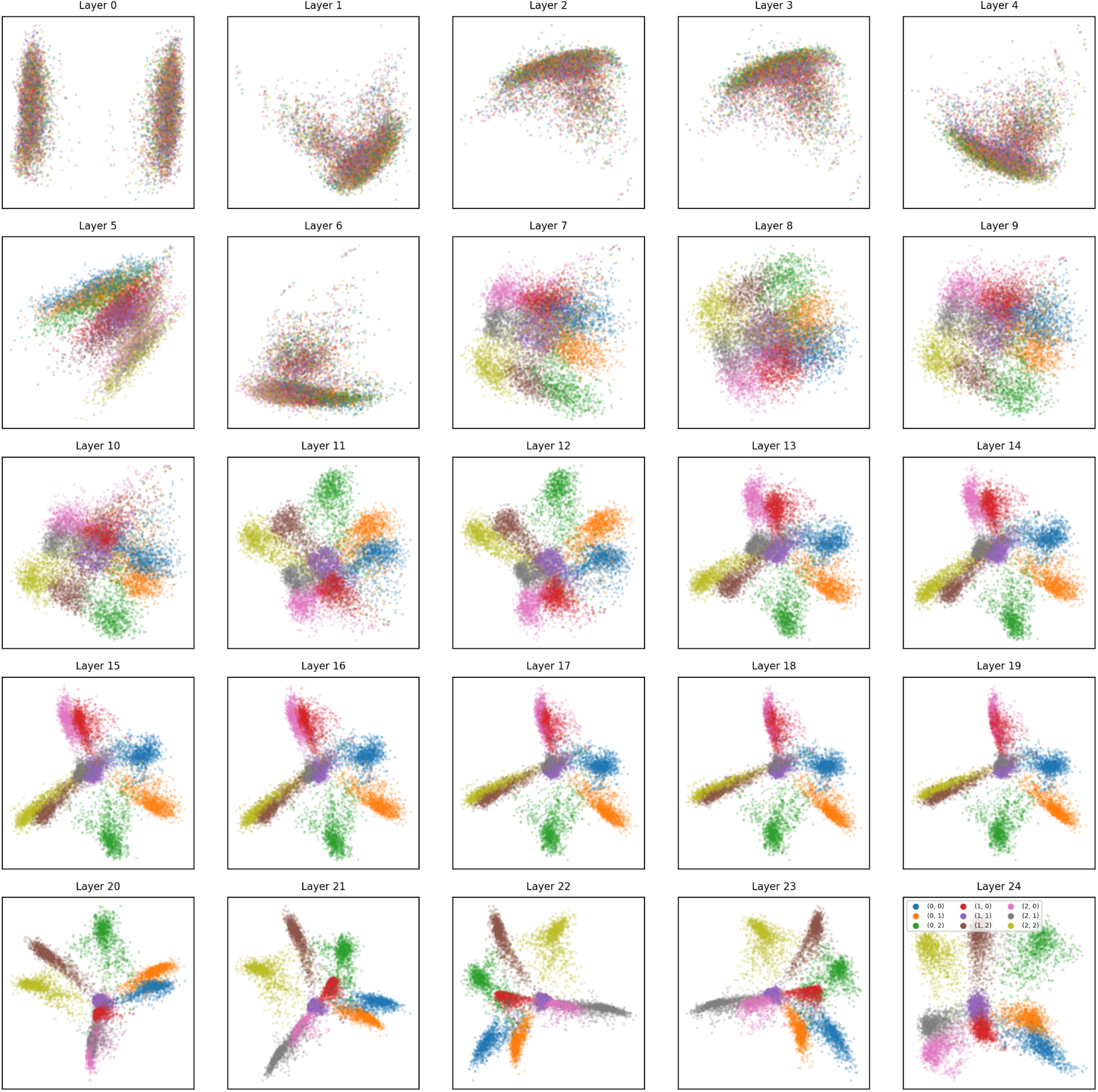
Principal component analysis of token representations in *all* layers of the spatial inference model in Section 2.3 (see Figure 7). Activations for each token of random walks in many different environments were extracted from a given layer of the model, to obtain an average vector for each location. Each subplot shows these representations projected into 2D with principal component analysis, colouring each location by its co-ordinates in the environment so that points with the same colour are different nodes which occupy the same position in the 3×3 grid.

**Table S1:** Narrative examples of retrieval-augmented generation with compression as a model of hippocampal-neocortical interaction, using Mistral-7B-Instruct-v0.2 plus xRAG weights to simulate neocortex. The ‘Original’ column gives the story ‘experienced’ by the system. The ‘Gist only’ column gives the reconstruction from just the xRAG vector, i.e. when the xRAG vector is retrieved from the hippocampus and used to condition the generative network. The ‘Stored detail’ column lists the most surprising detail given the xRAG vector. Specifically, all phrases are extracted from the text, then the perplexity of each phrase given the xRAG vector is calculated. In other words, for each phrase, the model is prompted with the xRAG vector and the phrase in question. Then the phrases are ranked in descending order of perplexity, with the top phrase used as the ‘Stored detail’. The text embedding (from which the xRAG vector is derived) and details for each memory are encoded together in the model hippocampus. The ‘Gist + detail’ column gives the reconstruction of the memory from this representation.

| Original | Gist only | Stored detail | Gist + detail |
| --- | --- | --- | --- |
| Sarah has a bad habit. All Sarah eats is candy and sweets. She starts to notice a sharp pain in her teeth. The dentist tells her she has 6 cavities! Sarah decides she needs to start eating healthier. | Sarah has a bad habit of only eating candy and not eating healthy foods. This is a problem because she is not getting the nutrients she needs to stay healthy. Her dentist told her that she has cavities and needs to start eating more fruits and vegetables . . . | The dentist tells her she has 6 cavities. | Sarah has a bad habit of eating only candy. The dentist tells her she has 6 cavities. Sarah’s bad habit of eating only candy has led to her having 6 cavities. The dentist told her this when she went for a check-up. Sarah needs to change her eating habits ... |
| Stan and his best friend have been friends for 8 years. They met when Stan was in first grade. They play games together, ate together, and watched movies together. Their favorite time of day is when Stan gets home from school. Stan’s best friend is always waiting for him with a wagging tail. | Stan and his best friend have been friends for 8 years. They met in kindergarten and have been inseparable ever since. They do everything together, from playing in the park to watching movies at home. They even have a special handshake they use to greet each other. Stan’s friend is always there for him, no matter what. . . . | Stan’s best friend is always waiting for him with a wagging tail. | Stan and his best friend have been friends for 8 years. They met when Stan was walking home from school and his friend was wagging his tail and barking at him. Stan approached the dog and petted him, and they have been friends ever since. They spend a lot of time together, playing, going for walks, and just hanging out. . . . |
| Gina prepared food for the family. She cooked lasagna, grilled chicken and garlic bread. As soon as we started to eat, her mom became nauseous. Afterwards, everyone started to feel queasy. They eventually threw away the food and ordered Chinese food. | Gina prepared food for the family. She cooked lasagna. However, she got sick and threw up. The family ate the lasagna, but they all got sick and threw up. They went to the hospital and were diagnosed with food poisoning. The food was later found to be contaminated with *E. coli*. | Her mom became nauseous. | Gina prepared food for the family. She cooked lasagna. However, her mom became nauseous and threw up after eating it. Gina then realized that she had used spoiled ground beef for the lasagna. The rest of the family also became sick after eating it. |

**Table S2:** Pearson correlations between story length and each LLM-derived attribute.

| Dataset | Concreteness | Richness in detail | Specificity |
| --- | --- | --- | --- |
| HIPPOCORPUS | $r = 0.019, p < 0.001$ | $r = 0.023, p < 0.001$ | $r = 0.023, p < 0.001$ |
| Simulated | $r = 0.279, p < 0.001$ | $r = 0.350, p < 0.001$ | $r = 0.240, p < 0.001$ |

**Table S3:** The results of t-tests on length-residualised attribute scores for HIPPOCORPUS stories.

| Attribute | Recalled vs. Retold | Recalled vs. Imagined | Retold vs. Imagined |
| --- | --- | --- | --- |
| Concreteness | $t = 4.823, p < 0.001$ | $t = 368.994, p < 0.001$ | $t = 6.445, p < 0.001$ |
| Richness in detail | $t = 5.467, p < 0.001$ | $t = 445.861, p < 0.001$ | $t = 7.770, p < 0.001$ |
| Specificity | $t = 7.772, p < 0.001$ | $t = 426.456, p < 0.001$ | $t = 5.292, p < 0.001$ |

**Table S4:** The results of paired t-tests on length-residualised attribute scores for simulated stories.

| Attribute | Original vs. Encoded | Original vs. Consolidated | Encoded vs. Consolidated |
| --- | --- | --- | --- |
| Concreteness | $t = 3.796, p < 0.001$ | $t = 5.850, p < 0.001$ | $t = 2.430, p = 0.015$ |
| Richness in detail | $t = 4.027, p < 0.001$ | $t = 6.710, p < 0.001$ | $t = 3.468, p < 0.001$ |
| Specificity | $t = 4.688, p < 0.001$ | $t = 6.581, p < 0.001$ | $t = 2.211, p = 0.028$ |

**Table S5:**
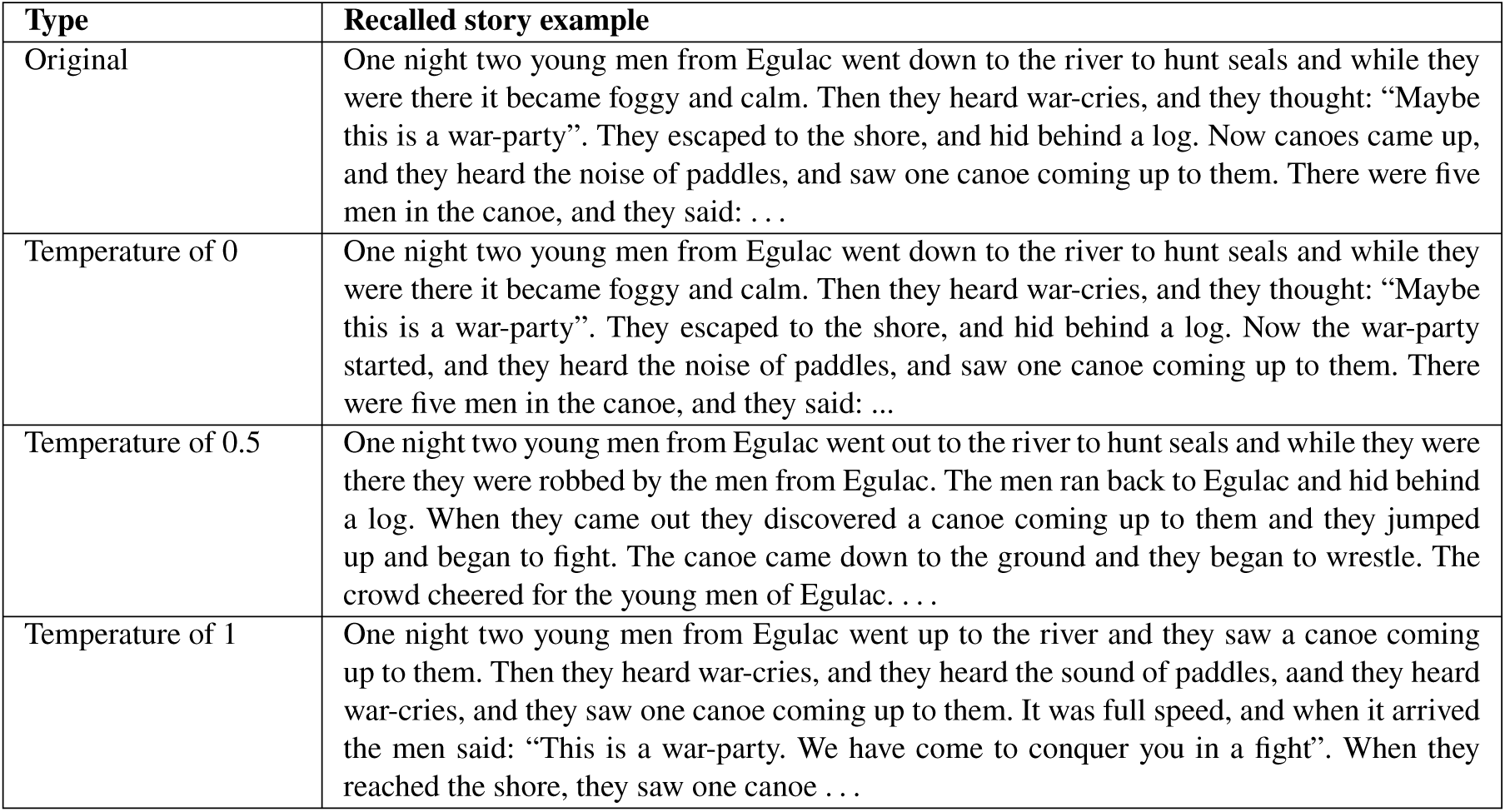
Recalled stories for different temperatures, for a model trained for five epochs with the ‘Sports’ category of Wikipedia data (*71*) as the background data distribution, and prompted with ‘One night two young men from Egulac’.

**Table S6:** Spatial task inference templates and their average accuracies.

| Inference template | Mean accuracy |
| --- | --- |
| { } EAST { } WEST { } | 1.0 |
| { } WEST { } EAST { } | 1.0 |
| { } NORTH { } SOUTH { } | 1.0 |
| { } SOUTH { } NORTH { } | 1.0 |
| { } EAST { } SOUTH { } WEST { } NORTH { } | 1.0 |
| { } SOUTH { } WEST { } NORTH { } EAST { } | 1.0 |
| { } WEST { } NORTH { } EAST { } SOUTH { } | 1.0 |
| { } NORTH { } EAST { } SOUTH { } WEST { } | 1.0 |
| { } EAST { } EAST { } NORTH { } WEST { } WEST { } SOUTH { } | 1.0 |
| { } NORTH { } NORTH { } WEST { } SOUTH { } SOUTH { } EAST { } | 1.0 |
| { } EAST { } EAST { } SOUTH { } SOUTH { } WEST { } WEST { } NORTH { } NORTH { } | 1.0 |
| { } SOUTH { } SOUTH { } WEST { } WEST { } NORTH { } NORTH { } EAST { } EAST { } | 1.0 |

**Table S7:**
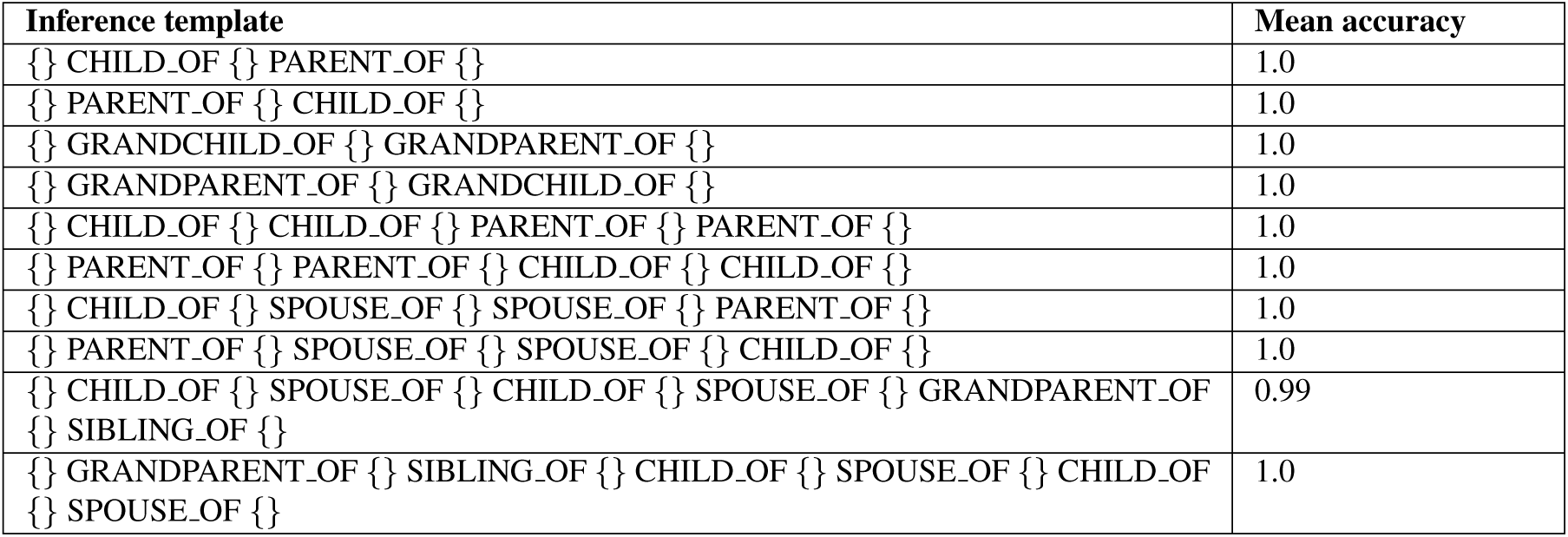
Family tree task inference templates and their average accuracies.

| Inference template | Mean accuracy |
| --- | --- |
| { } CHILD_OF { } PARENT_OF { } | 1.0 |
| { } PARENT_OF { } CHILD_OF { } | 1.0 |
| { } GRANDCHILD_OF { } GRANDPARENT_OF { } | 1.0 |
| { } GRANDPARENT_OF { } GRANDCHILD_OF { } | 1.0 |
| { } CHILD_OF { } CHILD_OF { } PARENT_OF { } PARENT_OF { } | 1.0 |
| { } PARENT_OF { } PARENT_OF { } CHILD_OF { } CHILD_OF { } | 1.0 |
| { } CHILD_OF { } SPOUSE_OF { } SPOUSE_OF { } PARENT_OF { } | 1.0 |
| { } PARENT_OF { } SPOUSE_OF { } SPOUSE_OF { } CHILD_OF { } | 1.0 |
| { } CHILD_OF { } SPOUSE_OF { } CHILD_OF { } SPOUSE_OF { } GRANDPARENT_OF { } SIBLING_OF { } | 0.99 |
| { } GRANDPARENT_OF { } SIBLING_OF { } CHILD_OF { } SPOUSE_OF { } CHILD_OF { } SPOUSE_OF { } | 1.0 |

